# A topologically conserved unstructured region helps positioning the evolutionarily conserved Prp40 WW domains to promote non-canonical intron splicing

**DOI:** 10.1101/2025.10.29.685309

**Authors:** Luh Tung, Chung-Shu Yeh, Hsin-Hung Lin, Hsin-Yue Tsai, Shang-Lin Chang, Amy Larson, Hsuan-Kai Wang, Fu-Lung Yeh, Chia-Ling Hsu, Cheng-Fu Kao, Jeffrey A. Pleiss, Wei-Hau Chang, Tien-Hsien Chang

## Abstract

Newly transcribed introns are immediately identified by early splicing factors that recognize the intron 5’ splice site (5’SS) and branch site (BS). In the budding yeast, these critical splice-site sequences are generally constrained, whereas degeneracy is the rule in higher eukaryotes. Yet, ∼40% of the yeast introns do diverge, to a certain degree, from the canonical sequences. Exactly how these non-canonical introns are recognized and spliced remains unknown. Here we show that the conserved Prp40 WW domains promote non-canonical intron splicing by enhancing stable U1 snRNP and BBP recruitments. AlphaFold predicts a topologically conserved unstructured region between Prp40 WW and FF domains. Alignment of the AlphaFold Prp40 structure with published U1 snRNP structure positions WW domains adjacent to 5’SS and Luc7, which is known to be critical for 5’SS recognition. Indeed, deletion of this unstructured region negatively impacts on splicing of the non-canonical 5’SS introns. Taken together, our results suggest that the conserved WW domains may have evolved to deal with the highly degenerate 5’SS and BS sequences in higher eukaryotes, so as to accommodate increased splicing complexity.

**Highlights:**

- The evolutionarily conserved Prp40 WW domains promote splicing of introns harboring non-canonical 5’ splice site or branch site.
- Prp40 WW domains enhance stable U1 snRNP and BBP recruitments to nascent transcripts containing non-canonical splice sites.
- A topologically conserved unstructured region between WW and FF domains helps to position Prp40 WW domains close to the 5’ splice site.
- The N-terminal WW domain sterically hinders conformational rearrangements required for efficient release of a BBP variant during spliceosome assembly.
- A reporter assay identified 13 non-canonical introns whose splicing, under various environmental conditions, depend on Prp40 WW domains.

## Introduction

Pre-mRNA splicing is a crucial process within the eukaryotic gene-expression pathway, responsible for removing introns and generating mature mRNA. This process is catalyzed by the spliceosome, a dynamic molecular machine that mediates intron excision and exon ligation through two sequential transesterification reactions. The spliceosome comprises five small nuclear ribonucleoproteins (snRNPs: U1, U2, U4, U5, and U6) and the NineTeen Complex (NTC), and it assembles stepwise on intron-containing transcripts^1,2^. Cis-acting sequences surrounding the intron 5’ splice site (5’SS), branch site (BS), and 3’ splice site (3’SS) are critical for RNA-RNA and RNA-protein interactions that mediate this stepwise spliceosome assembly^3,4^. Upon transcription, the 5′SS of a nascent RNA is initially recognized by U1 snRNP, followed by the recruitment of U2 snRNP to the BS, forming the pre-catalytic spliceosome^5,6^. In hemiascomycetous yeasts, most 5′SS and BS cis-elements are conserved^7^. For example, in *Saccharomyces cerevisiae*, 75% (233/314) of 5′SS sequences conform to the consensus GUAUGU, and 80% (253/314) of BS sequences are UACUAAC (Fig 1A). Nonetheless, a substantial fraction of introns (∼40%) contain noncanonical 5′SS and/or BS sequences (Fig. 1B). The mechanism by which these noncanonical introns are efficiently spliced remains poorly understood.

**Fig. 1.**
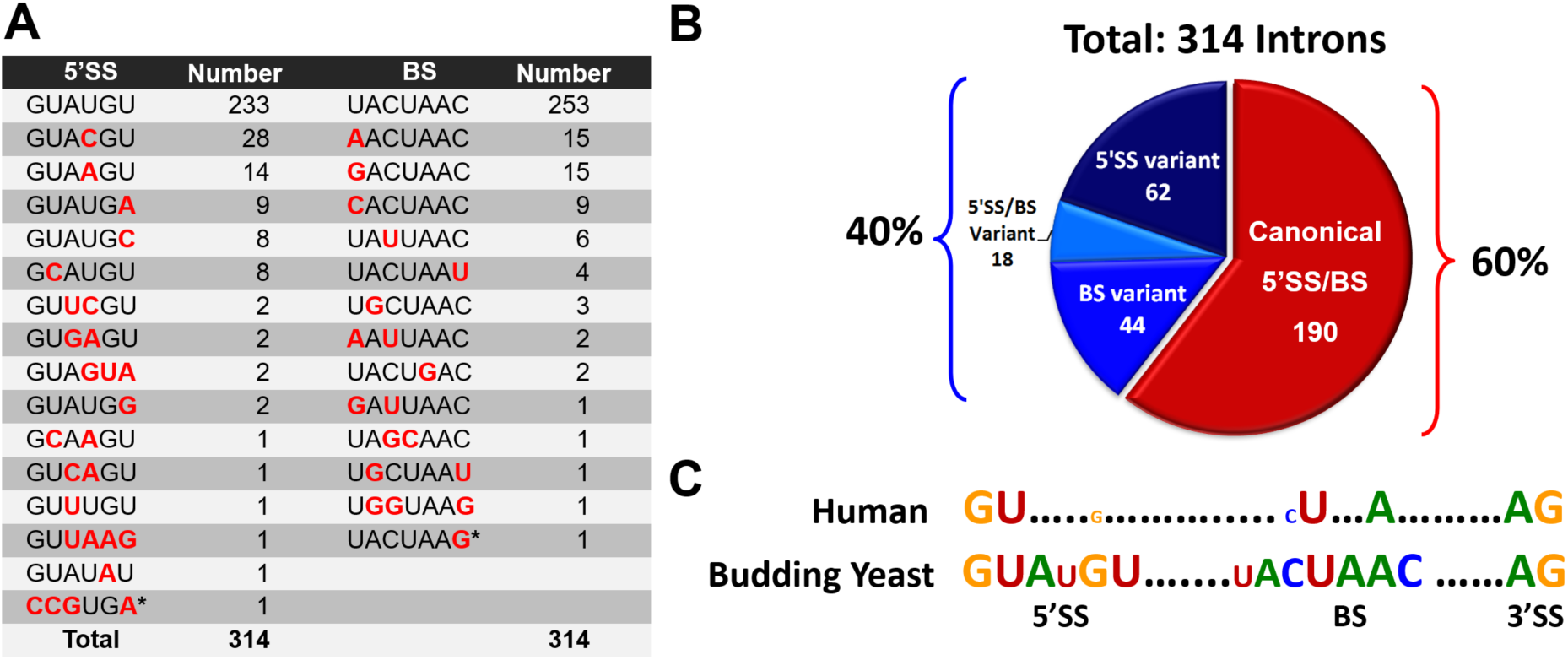
Compiled yeast 5’SS and BS sequences. (A) Splice-site sequences and their frequencies in the budding yeast. Nucleotides deviating from the canonical sequences are highlighted in red. Asterisks denote the 5’SS and BS of the *HAC1* intron, which undergoes spliceosome-independent splicing mediated by Ire1 and tRNA ligase. (B) Distribution of canonical and variant splice-site sequences. (C) Human intron 5’SS and BS sequences are far more degenerate in comparison to those in the budding yeast.

Prp40 is an essential component of U1 snRNP in ascomycete yeast^8,9^ and has orthologs in *Caenorhabditis elegans*, *Drosophila melanogaster*, *Homo sapiens*, and *Arabidopsis thaliana*^10^ (S1). Prp40 possesses evolutionarily conserved domains, including two tandem WW repeats at its N-terminus, followed by six FF domains^11–13^. The WW domain is about 23 amino-acid long, with a tryptophan at both ends of the domain (S2). It was initially proposed that Prp40’s WW domain interacts with the branch-site binding protein’s (BBP) PPVY region, potentially bringing 5’SS and BS into close proximity^11^. Structural analysis revealed that the WW domain forms three beta sheets connected by two loops^12^. WW domain-containing proteins are found exclusively in eukaryotes and have been implicated in regulating expression, localization, and stability of their targets in various biological processes^13^, including transcription^14,15^, apoptosis^14,15^, splicing^8,11,16,17^, ubiquitylation^18^, tumorigenesis^19,20^, and many human neurological diseases^13,21–26^.

A prior study suggested that the yeast Prp40 WW domains are dispensable for splicing *in vitro*, commitment-complex formation, and co-transcriptional spliceosome assembly^27^. This conclusion appeared to contradict the high degree of conservation of the Prp40 WW domains across eukaryotes. In this study, we re-examined this apparent paradox from a new perspective. We demonstrate that Prp40’s WW domains are, in fact, critical for splicing non-canonical 5’SS and BS-containing introns. Current spliceosome structures^28,29^ do not provide information regarding the precise location of WW domains within the spliceosome or their spatial relationship to the 5’SS and BS. Leveraging the predictive power of AlphaFold^30^, and combining this with existing spliceosome structures, we identified a topologically conserved unstructured region between the WW and FF domains of Prp40 (S1A). This region may allow the WW domains to approach the 5′SS and interact with Luc7^31^, a known factor in 5′SS recognition. Supporting this model, deletion of the unstructured linker region impairs the splicing efficiency of introns with noncanonical 5′SS. We propose that the conserved WW domains of Prp40 have evolved to support the splicing of noncanonical introns and may play an increasingly important role in higher eukaryotes, where splice-site sequences are markedly degenerate.

## Results

### Prp40 WW domains genetically interact with a group of splicing factors

The evolutionarily conserved Prp40 WW domains were previously proposed to mediate the interaction between the U1 snRNP and BBP during early spliceosome assembly^11,12,32^. Yet, yeast strains lacking both WW domains exhibited no apparent growth and splicing defects^27^. To resolve this discrepancy, we constructed three isogenic strains: *prp40-ΔWW1* (lacking the first WW domain), *prp40-ΔWW2* (lacking the second WW domain), and *prp40-ΔN* (lacking both WW domains) (S2A and S2B). Consistent with previous findings^27^, we found no discernible growth difference between the wild-type strain and the three mutants (*prp40-ΔWW1, prp40-ΔWW2, and prp40-ΔN*) on solid medium (S2B) or in a more sensitive competitive fitness assay in liquid medium at 30°C (see Materials and Methods) (S2C).

To genetically identify genes that may functionally interact with the WW domains, we performed a genome-wide synthetic genetic array (SGA) screen^33,34^. We crossed the wild-type and the three mutant *prp40* strains with a panel of approximately 4,300 haploid nonessential-gene deletion strains (see Materials and Methods). This screen revealed nine gene deletions that, when combined with the mutant *prp40* alleles, resulted in a statistically significant reduction in fitness (Table 1). Notably, *MUD2*, *LEA1*, *BRR1*, and *LSM6* among these nine genes are known to be involved in splicing. Both Mud2 and BBP interact with Prp40 genetically and physically^11^ and form a heterodimer that binds to the intron branch site region^35^, a region subsequently occupied by U2 snRNP. A recent pre-spliceosome structure^36^ showed that Lea1, a U2-snRNP component, directly contacts Prp39, a protein of the U1 snRNP^37^. Brr1, the third candidate, genetically interacts with Cbc2, the small subunit of the nuclear cap-binding complex^38^, which also genetically interacts with Mud2 and BBP^38^. Collectively, these data strongly suggest that the Prp40 WW domains play a role in the formation and/or stability of commitment complexes, which include Prp40, Mud2, BBP, and Cbc2.

**Table 1.**
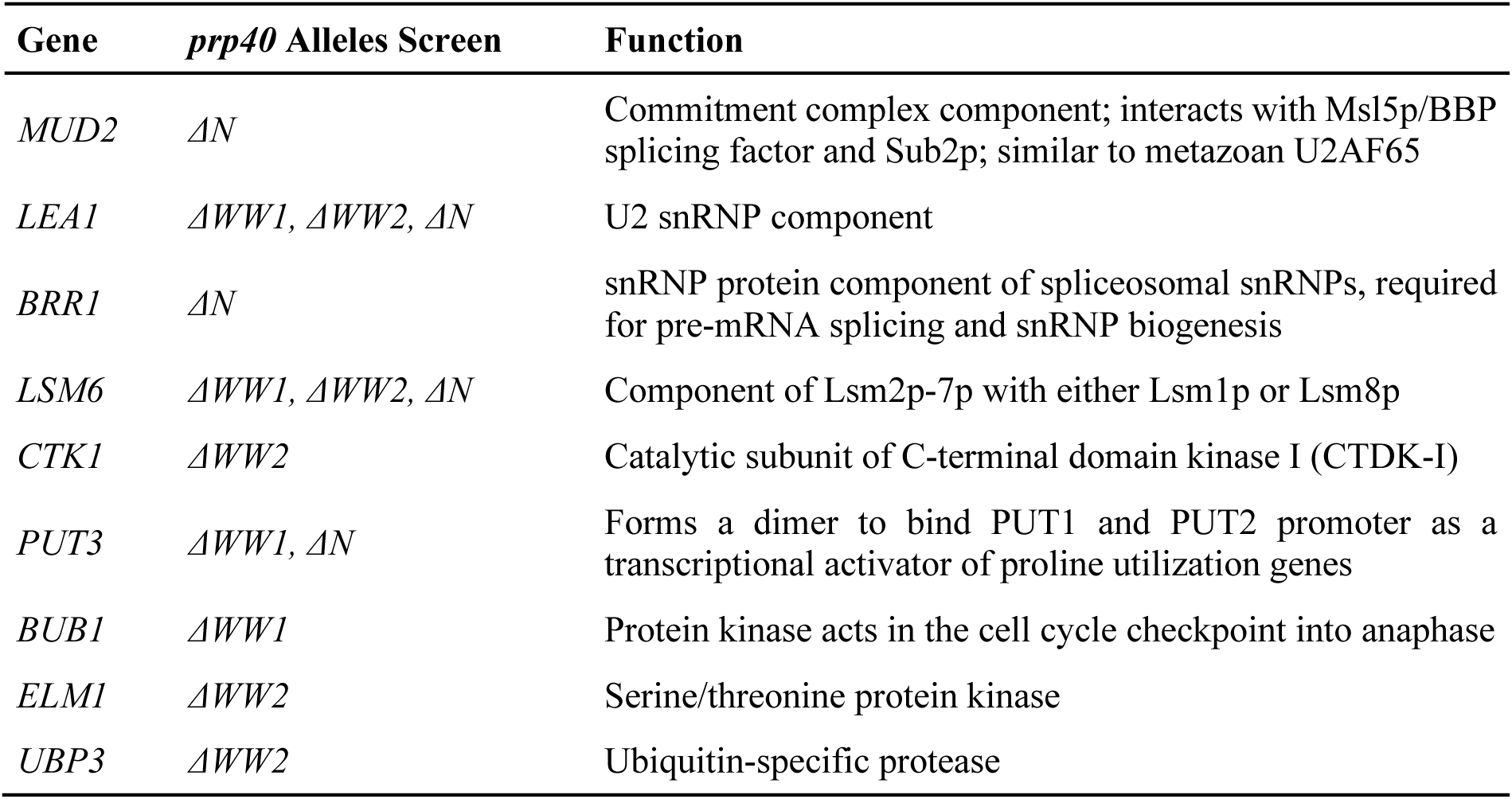
SGA screen result.

| Gene | *prp40* Alleles Screen | Function |
| --- | --- | --- |
| *MUD2* | $\Delta N$ | Commitment complex component; interacts with Msl5p/BBP splicing factor and Sub2p; similar to metazoan U2AF65 |
| *LEA1* | $\Delta WW1, \Delta WW2, \Delta N$ | U2 snRNP component |
| *BRR1* | $\Delta N$ | snRNP protein component of spliceosomal snRNPs, required for pre-mRNA splicing and snRNP biogenesis |
| *LSM6* | $\Delta WW1, \Delta WW2, \Delta N$ | Component of Lsm2p-7p with either Lsm1p or Lsm8p |
| *CTK1* | $\Delta WW2$ | Catalytic subunit of C-terminal domain kinase I (CTDK-I) |
| *PUT3* | $\Delta WW1, \Delta N$ | Forms a dimer to bind PUT1 and PUT2 promoter as a transcriptional activator of proline utilization genes |
| *BUB1* | $\Delta WW1$ | Protein kinase acts in the cell cycle checkpoint into anaphase |
| *ELM1* | $\Delta WW2$ | Serine/threonine protein kinase |
| *UBP3* | $\Delta WW2$ | Ubiquitin-specific protease |

### Prp40 WW domains are required for efficient splicing of introns with non-canonical 5’SS or BS sequences

To define the role of Prp40 WW domains in splicing, we utilized a splicing-sensitive microarray^39,40^ to evaluate the efficiency of splicing for all known yeast introns (S3A). We focused on detecting reductions in spliced transcripts within the *prp40-ΔN* background (see Materials and Methods). The impact observed was considerably less severe than that seen in the *prp2* or *prp8* mutants^39,40^ (S3A). Nevertheless, we identified eleven candidate genes (S3B). Upon closer examination of their intron sequences, nine shared a common characteristic: their 5’SS or BS sequences deviated from the canonical GUAUGU (5’SS) and UACUAAC (BS) consensus sequences (S3B). This observation led to the hypothesis that the splicing of these nine introns is particularly sensitive to the absence of Prp40 WW domains. To validate this finding, we directly measured splicing efficiency via RT-qPCR in both *prp40-ΔN* and *PRP40* genetic backgrounds (Fig. 2A). Five other genes containing canonical introns served as controls. As predicted, significantly higher levels of unspliced transcripts were detected in the *prp40-ΔN* background for the nine candidates compared to the controls (Fig. 2A). A particularly striking example is the *RPL7B* gene, which contains two introns: the first intron (RPL7B1) has a non-canonical branch site (U<u>G</u>CUAAC), while the second intron (RPL7B2) has the typical UACUAAC. Consistent with our hypothesis, the splicing of RPL7B1 was more significantly affected by the deletion of the Prp40 WW domains than that of RPL7B2 (Fig. 2A). These findings suggest that deletion of Prp40 WW domains impairs the splicing of a subset of introns containing non-canonical *cis*-acting elements at either the 5’SS or the BS.

**Fig. 2.**
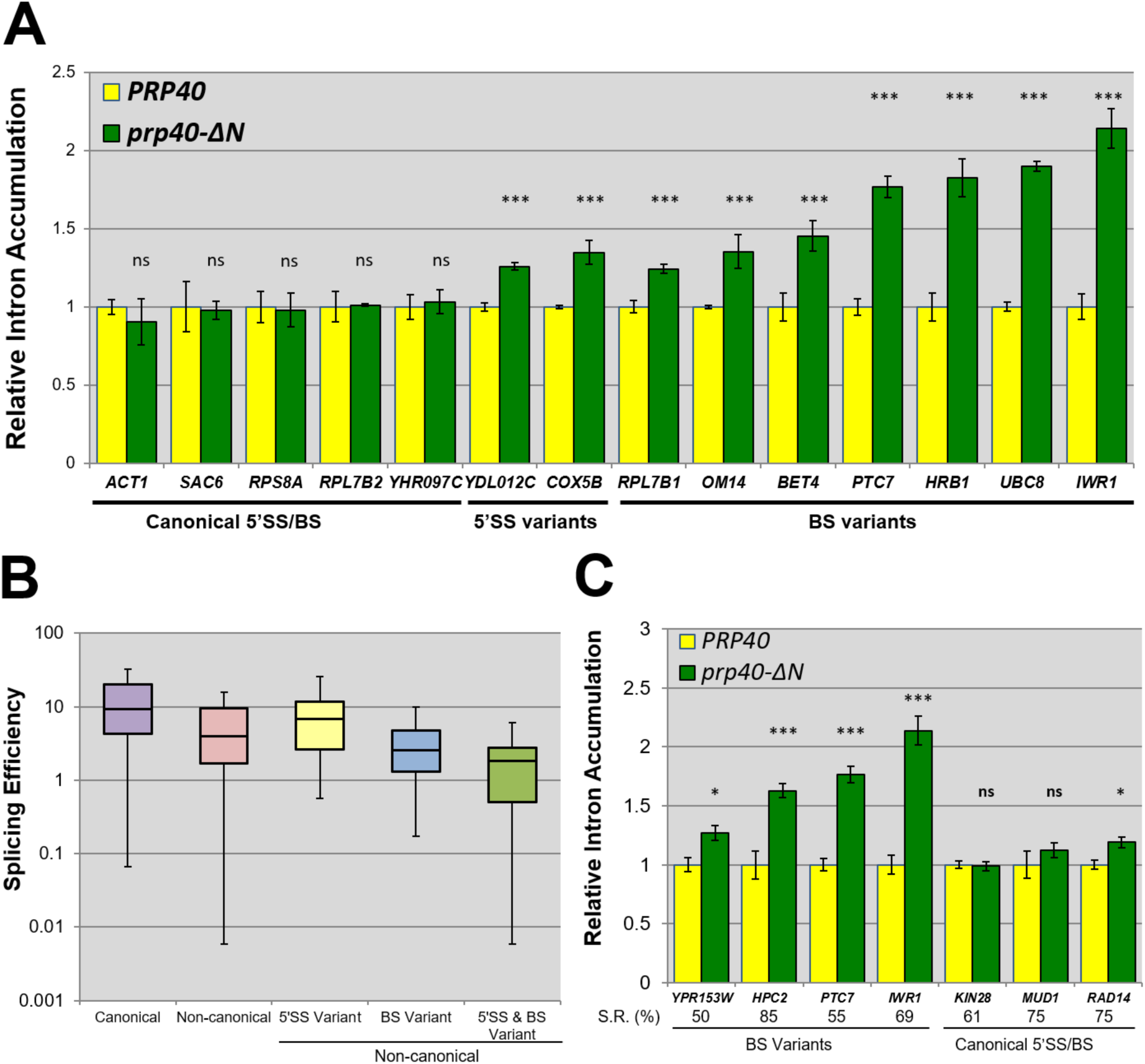
Splicing efficiencies of various introns in *PRP40* and *prp40-ΔN* yeast strains. (A) Relative intron accumulation in *PRP40* and *prp40-ΔN* backgrounds. Intron accumulation was quantified for transcripts containing canonical 5′SS/BS, variant 5′SS, or variant BP. Splicing efficiency was calculated based on intron accumulation. Increased levels of introns indicate decreased splicing efficiency. The ratio of intron to spliced mRNA was determined by RT-qPCR for each transcript and normalized to the corresponding value in the *PRP40* background to generate relative intron accumulation values (Y-axis). *N* = 4 biological repeats; error bars, ±SEM; \**P* < 0.05, \*\**P* < 0.01, \*\*\**P* < 0.001, and ns, no statistical significance. (B) Splicing efficiency of canonical and non-canonical introns calculated from RNA-seq data. RNA-seq was done in the wild-type genetic background, the data of which were first grouped into two categories, canonical introns and non-canonical introns. The non-canonical data were further categorized into three separate groups, introns containing 5’SS variants, BS variants, as well as both 5’SS and BS variants. (C) Inefficiently spliced introns were examined for their splicing efficiencies as described in (A). Literature^41^-reported splicing rates (S.R. [%]) in the *PRP40* background are indicated below the X-axis. *N* = 3 biological repeats; error bars, ±SEM; \**P* < 0.05, \*\**P* < 0.01, \*\*\**P* < 0.001, and ns, no statistical significance.

Having established this specificity, we next asked whether non-canonical introns are intrinsically less efficiently spliced, which could underlie their sensitivity to WW-domain loss. RNA-seq analysis of all pre-mRNA splicing events in wild-type *PRP40* cells revealed that introns with non-canonical 5′SS and/or BS sequences indeed exhibit reduced splicing efficiency relative to canonical introns (Fig. 2B). Among these, introns carrying both non-canonical 5′SS and BS were the least efficiently spliced, followed by those with a non-canonical BS, whereas non-canonical 5′SS alone had the mildest effect (Fig. 2B). Thus, non-canonical introns represent intrinsically less efficiently spliced substrates *in vivo*.

To determine whether this intrinsic inefficiency alone accounts for their dependence on the Prp40 WW domains, we examined canonical introns that are also inefficiently spliced. Two of the candidate genes identified through our microarray analysis, *IWR1* (69.2% spliced) and *PTC7* (55.3%), are known to exhibit low splicing efficiency *in vivo*^41^. This observation prompted us to investigate whether the sensitivity to loss of Prp40 WW domains could be attributed to the inherent splicing inefficiency of these genes, rather than the presence of non-canonical intron *cis*-acting elements. If the observed effect were solely due to poor intrinsic splicing efficiency, we would expect that all genes with low splicing efficiency^41^, regardless of their intron type, would show further impairment in the *prp40*-*ΔN* background. To test this hypothesis, we performed RT-qPCR analysis on five genes known to be inefficiently spliced (Fig. 2C). These included *KIN28* (61%), *MUD1* (75%), and *RAD14* (75%), which contain canonical introns, and *HPC2* (85%) and *YPR153W* (50%) which contain non-canonical BS introns. Our results demonstrated that the splicing of *HPC2* and *YPR153W* was significantly reduced in the *prp40*-*ΔN* background, while the splicing of *KIN28, MUD1,* and *RAD14* remained largely unaffected. These findings support the conclusion that Prp40 WW domains play a specific role in facilitating the splicing of non-canonical introns, independent of their intrinsic splicing efficiency *in vivo*.

### Prp40 WW domains compensate for non-canonical intron splicing *in vivo*

To comprehensively assess the impact of the loss of Prp40 WW domains on non-canonical intron splicing, we surveyed the Ares Intron Database (http://intron.ucsc.edu/yeast4.1/)^42^ and Saccharomyces Genome Database (http://www.yeastgenome.org)^43^ to characterize intron *cis*-information in yeast. The most prevalent 5’SS and BS sequences were identified as GUAUGU (233/314; 75%) and UACUAAC (253/314; 80%), respectively (Fig. 1A). However, a range of non-canonical sequences, deviating by up to four mismatches, were also observed. Approximately 5.7% (18/314) of surveyed introns displayed non-consensus sequences in both the 5’SS and BS regions (Fig. 1B). While yeast introns are generally more constrained than their human counterparts (Fig. 1C), a substantial proportion (almost 40%) deviate from the prevalent architecture (Figs. 1A and 1B). Based on this observation, we hypothesized that the conservation of Prp40 WW domains may compensate for the presumed suboptimal splicing efficiency of these non-canonical introns.

To evaluate the effect of specific 5’SS and/or BS variants (Fig. 1A) on splicing efficiency *in vivo*, we utilized an established *ACT1/CUP1* splicing reporter system^44^. This system fuses a fragment of the *ACT1* gene (including exon 1, intron, and a portion of exon 2) with the *CUP1* open reading frame. Introduction of this reporter construct on a plasmid allows us to measure splicing efficiency based on cell growth on plates with varying CuSO_4_ concentrations^44^. We generated 31 reporter constructs, incorporating 28 naturally occurring variants and three previously characterized artificial variants^27^. These constructs were introduced into yeast strains with both *PRP40* and *prp40*-*ΔN* backgrounds, and growth phenotypes were assessed on plates with varying CuSO_4_ concentrations. A complete dataset of results is provided in S4. Figures 3A and 3B illustrate two examples of this analysis. In the *prp40*-*ΔN* background, the 5’SS variant GU<u>GA</u>GU resulted in cell lethality at all tested temperatures on plates with 0.1 mM CuSO_4_ (Fig. 3A). Conversely, robust cell growth was observed in the *PRP40*, *prp40*-*ΔWW1*, and *prp40*-*ΔWW2* backgrounds at both 18°C and 30°C. Similarly, the BS variant <u>G</u>A<u>U</u>UAAC resulted in a severe growth phenotype specifically in the *prp40*-*ΔN* background when grown on plates with 0.5 mM CuSO_4_ (Fig. 3B). Some variants, such as GUAUG<u>C</u>, GUA<u>GUA</u> (5’SS), and UACUAA<u>U</u> (BS), exhibited a slow growth phenotype at 18°C (S4). A subset of variants either caused cell death (3/17 for 5’SS and 1/14 for BS), which could be due to the heterologous nature of the *ACT1/CUP1* reporter system.

**Fig. 3.**
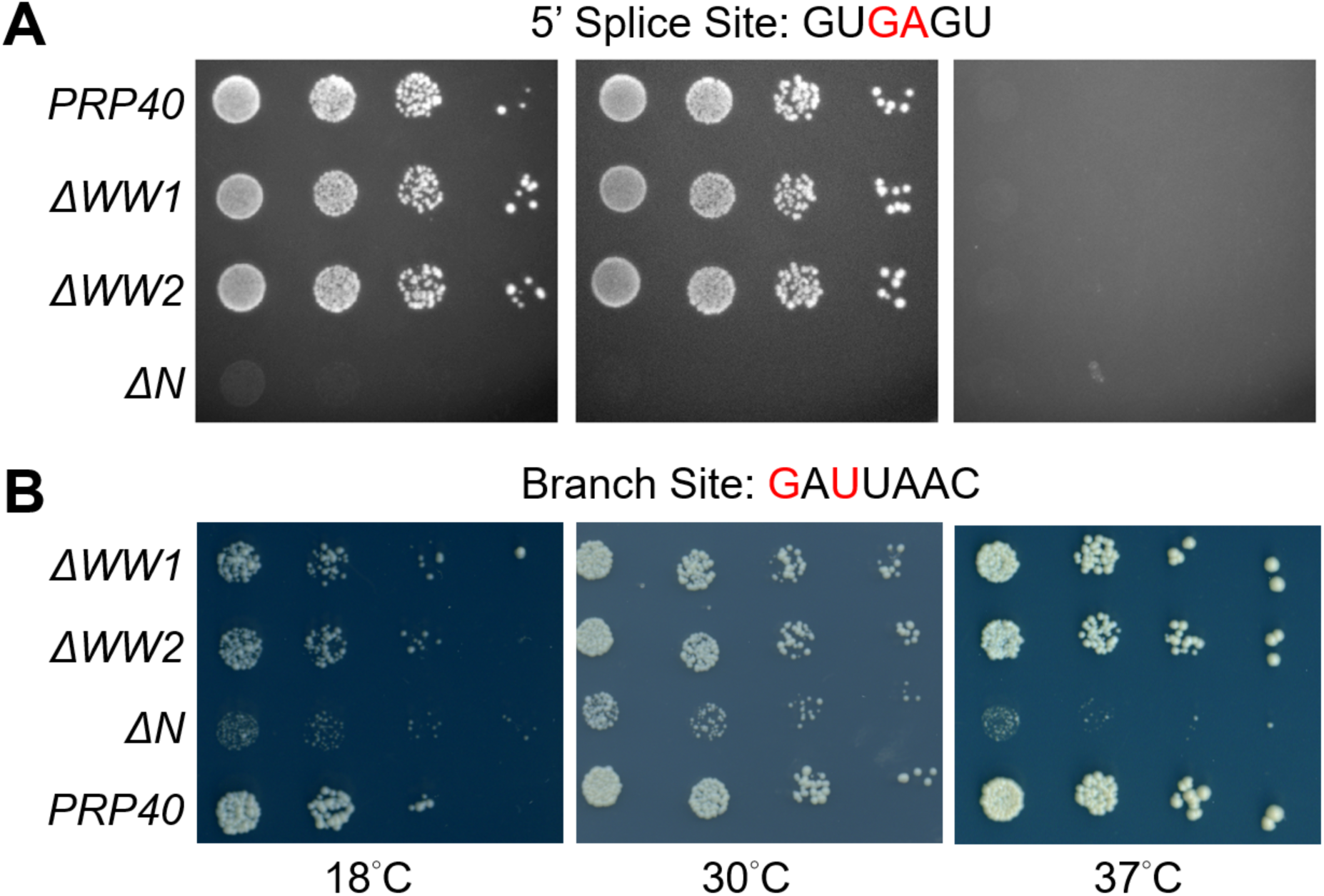
Growth phenotypes of *PRP40* and *prp40* mutant strains containing a copper-sensitive splicing reporter with noncanonical splice-site sequences. (A) Phenotypes of strains containing a copper-sensitive splicing reporter with 5’SS variant GU<u>GA</u>GU (non-canonical nucleotides are shown in red). Strains examined are in the *PRP40*, *ΔWW1* (deletion of the first WW domain), *ΔWW2* (deletion of the second WW domain), and *ΔN* (deletion of both WW domains) genetic backgrounds. Yeast strains of the corresponding genotypes were serially diluted and spotted onto copper-containing plates and incubated at the specified temperatures. (B) Phenotypes of strains harboring a copper-sensitive splicing reporter with BS variant <u>G</u>A<u>U</u>UAAC.

Through this analysis, we identified 13 additional intron variants (S4) whose *in vivo* splicing efficiency is affected by the absence of Prp40 WW domains. Interestingly, inspection of base-pairing interactions between splice sites and snRNAs (5’SS/U1 snRNA and BS/U2 snRNA) revealed no clear correlation with the role of Prp40 WW domains in splicing (S5).

### Prp40 WW domains stabilize U1 snRNP and BBP recruitment to non-canonical introns

Previous research have demonstrated that Prp40 physically interacts with BBP^11^ and its WW domains establish structural contacts with BBP’s proline-rich PPVY peptide^12,32^. Based on these findings, we hypothesized that physical contacts between U1 snRNP and BBP, facilitated by Prp40 WW domains, are critical for forming stable early splicing complexes, such as CC2, on introns containing non-canonical 5’SS or BS sequences. These non-canonical sequences are considered sub-optimal spliceosome substrates. Conversely, we anticipated that these contacts would be less critical for introns containing canonical sequences. This hypothesis regarding canonical introns is supported by a previous study^27^, which observed no splicing defects when using canonical or non-functional introns inactivated at the 5’ SS or BS. To further investigate the mechanistic role of the Prp40 WW domains, we performed chromatin immunoprecipitation (ChIP) to monitor the recruitment of U1 snRNP and BBP to three intron-containing genes: *ACT1* (containing canonical 5’SS and BS sequences), *HPC2* (containing a non-canonical branch site <u>G</u>A<u>U</u>UAAC), and *RPL20A* (containing a non-canonical 5’SS GU<u>GA</u>GU). Our results showed that recruitments of both U1 snRNP and BBP was less efficient for non-canonical *HPC2* and *RPL20A* introns in the *prp40-ΔN* background compared to the wild-type *PRP40* background, while recruitment remained similar for *ACT1* (Fig. 4A).

**Fig. 4.**
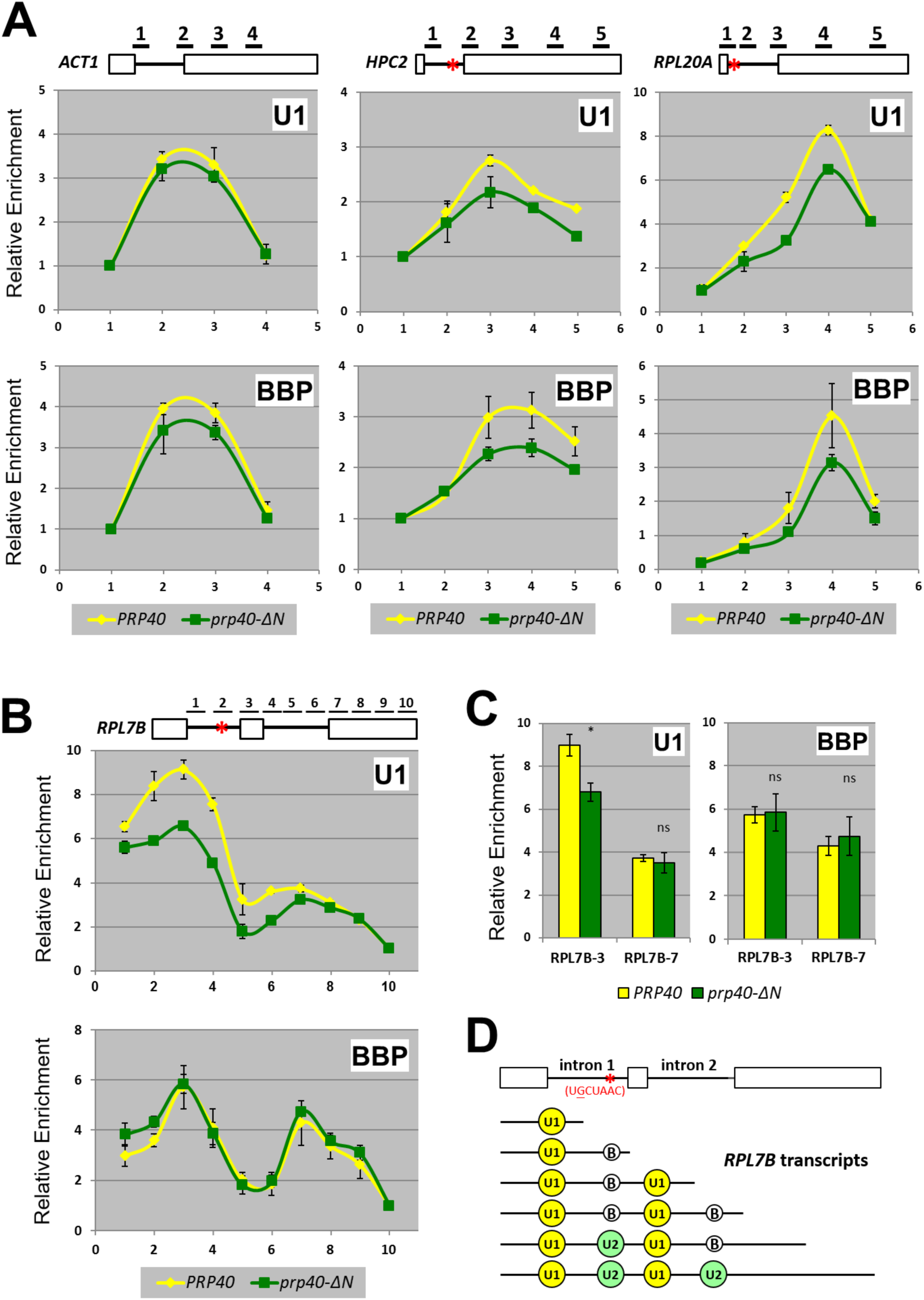
Co-transcriptional association of U1 snRNP and BBP with introns containing canonical or non-canonical splice sites. (A) Chromatin immunoprecipitation (ChIP) was used to assess co-transcriptional association of U1 snRNP and BBP with the introns of *ACT1*, *HPC2* (BS variant), and *RPL20A* (5’SS variant) in the *PRP40* (yellow line) or the *prp40-ΔN* (green line) genetic background. U1C-TAP (U1 snRNP) and MSL5-TAP (BBP) were used as tagged proteins for the purpose of immunoprecipitation. In the top diagram, exons are depicted as boxes, introns as lines, and red asterisks mark splice-site variants. DNA enrichment (Y-axis) was analyzed using primer sets denoted by numbers in the top diagram and below the X-axis. All data were normalized to the value obtained from the first primer set. *N* = 3 biological repeats; error bars, ±SEM. (B) Association of the U1 snRNP and BBP to the *RPL7B* introns*. RPL7B* has two introns, with the first intron harboring a non-canonical BS. *N* = 3 biological repeats; error bars, ±SEM. (C) Quantitative representation of peak data points from primer sets 3 and 7 from (B). RPL7B-3, primer set 3; RPL7B-7, primer set 7. *N* = 3 biological repeats; error bars are ±SEM; \**P* < 0.05, \*\**P* < 0.01, \*\*\**P* < 0.001, and ns, no statistical significance. (D) Conceptual presentation of the co-transcriptional association of U1 snRNP and BBP with *RPL7B*’s two introns.

We further investigated U1 snRNP and BBP recruitment by performing the same analysis using a series of ten oligo pairs covering the length of *RPL7B* (Fig. 4B). *RPL7B* contains two introns: intron 1 (409 nucleotides) contains a non-canonical BS sequence (U<u>G</u>CUAAC), and intron 2 (410 nucleotides) contains canonical sequences at both sites (Fig. 4B). While only intron 1 is sensitive to the loss of Prp40 WW domains (Fig. 2A), our analysis revealed that both U1 snRNP and BBP were recruited onto the *RPL7B* transcript in two distinct waves, peaking at oligo pairs 3 and 7 (Fig. 4B), respectively, consistent with the presence of two introns. The level of U1 snRNP recruitment was higher during the first wave than the second, reflecting the later appearance of the intron-2-containing region during transcription (Fig. 4B–D). Importantly, we observed a lower level of co-transcriptionally recruited U1 snRNP on intron 1 in the *prp40-ΔN* background compared to the *PRP40* background (Fig. 4B–D), while the recruitment levels in the canonical intron 2 were similar in both backgrounds (Fig. 4B–D).

Although the levels of U1 snRNP recruitment clearly differed between intron 1 and intron 2, BBP levels were similar for both introns. This can be explained by the fact that a fraction of BBP is replaced by U2 snRNP during earlier spliceosomal assembly at intron 1 compared to intron 2 (Fig. 4D). Counterintuitively, we observed similar BBP levels at the non-canonical intron 1, which harbors a non-canonical BS sequence (U<u>G</u>CUAAC), in both genetic backgrounds (Fig. 4B, bottom). We suspect that this could be due to the non-canonical BS sequence primarily affects the physical arrangement of BBP within the spliceosome, rather than directly affecting BBP binding to the branch site. And that may in turn impair the stable association of U1-snRNP within the spliceosome. Taken together, our data strongly support the idea that Prp40 WW domains promote the stable association of U1 snRNP and BBP during spliceosomal formation on introns containing non-canonical *cis*-acting elements.

### Prp40 WW single-domain deletion rescues splicing defect caused by specific BBP variants

To investigate the functional relationship between BBP (encoded by the *MSL5* gene) and the Prp40 WW domains, we engineered two mutant *msl5* alleles, predicted to compromise BBP’s physical interaction with the intron branch-site region based on published structural data^12,32,38^. These mutant alleles, *msl5*-S194P and *msl5*-V195D, exhibited a cold-sensitive phenotype at 16°C when expressed in an otherwise wild-type *PRP40* background (Fig. 5A; left panel; rows 1 and 5). As genetically anticipated, both *msl5*-S194P and *msl5*-V195D exacerbated the cold-sensitive phenotype when combined with the *prp40-ΔN* allele (Fig. 5A; left panel; rows 1 vs. 4 and rows 5 vs. 8). The *msl5-*S194P *prp40-ΔN* strain’s fitness reduction cannot be easily detected on plate at 30°C, but can be quantified by the quantitative fitness assay (Fig. 5B). In contrast, *prp40-ΔWW1* and *prp40-ΔWW2* alleles displayed no growth phenotype at 30°C (Fig. 5B). Interestingly, contrary to our initial prediction, both *prp40-ΔWW1* and *prp40-ΔWW2* alleles rescued, rather than worsened, the growth phenotype of *msl5-*S194P at 16°C (Fig. 5A; left panel; rows 2, 3, 6, and 7).

**Fig. 5.**
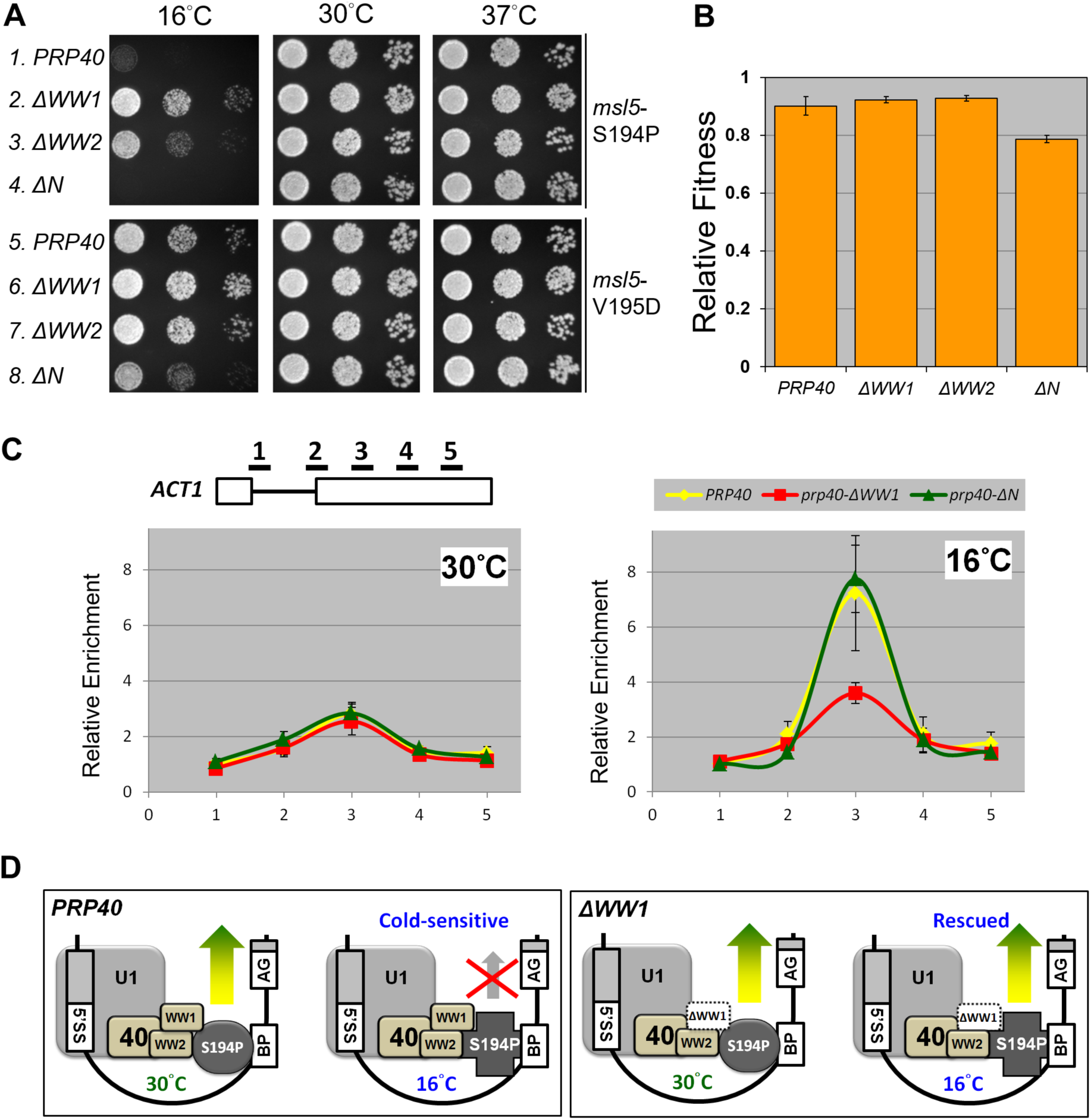
Deletion of a single WW domain in Prp40 rescues the cold-sensitive phenotype of two BBP mutant strains. (A) The *msl5*-S194P and *msl5*-V195D mutant strains exhibit growth defects at 16°C in the wild-type *PRP40* background (rows 1 and 5; left panel). Deletion of a single WW domain—either the first (*ΔWW1*) or the second (*ΔWW2*)—partially rescues the cold-sensitive phenotype (rows 2 and 3; 6 and 7; left panel). Deletion of both WW domains (*ΔN*, rows 4 and 8; left panel) exacerbates the growth defect at 16°C. (B) Competitive fitness assay of the wild-type and mutant *PRP40* strains in the *msl5*-S194P genetic background. Even at 30°C, deletion of Prp40’s two WW domains (*ΔN*) compromises cell fitness, which cannot be visualized by spotting test in (A) (row 4, middle panel). (C) Co-transcriptional association of the BBP-S194P variant with the *ACT1* intron was analyzed by ChIP at 30°C and 16°C. ChIP signals are presented for strains of wild-type *PRP40* (yellow), *prp40-ΔWW1* (red), or *prp40-ΔN* (green) to illustrate the impact of WW domain deletion on BBP recruitment. *N* = 3 biological repeats; error bars are ±SEM. (D) A proposed model illustrating how a single WW domain deletion may rescue the cold-sensitive phenotype of the mutant *msl5*-S194P strain. In the *msl5*-S194P genetic background, the presence of both WW domains appears to impede BBP’s release as shown in (C). The absence of either of the two WW domains provides a pathway for BBP to dissociate, thus facilitating the subsequent U2-snRNP binding to the branch site at 16°C.

To explain this observation, we investigated the co-transcriptional recruitment of BBP-S194P to the *ACT1* transcript in both the *PRP40* and *prp40-ΔWW1* backgrounds by ChIP analysis (Fig. 5C). We focused on BBP-S194P, because the cold-sensitive phenotype of *msl5*-S194P was more prominent than that of the *msl5*-V195D (Fig. 5A; left panel; rows 1 vs. 5). Our findings revealed that in the *PRP40* background at 16°C, the association of BBP-S194P with the transcript (Fig. 5C; right panel; yellow line) was significantly higher (∼2.5-fold) compared to that of the 30°C (Fig. 5C; left panel; yellow line), raising the possibility that BBP-S194P may not be efficiently released from the altered spliceosomal environment.

Notably, the *prp40-ΔWW1* allele appeared to rescue the biochemical phenotype of *msl5*-S194P (Fig. 5C; right panel; red line), consistent with the genetic suppression observed for both *msl5-*S194P and *msl5-*V195D in the absence of the WW1 domain (Fig. 5A; left panel). This finding suggests that the loss of the Prp40 WW1 domain may have created a spliceosomal conformation that facilitated the release of BBP-S194P (Fig. 5D). Collectively, our genetic and biochemical studies provided strong support for a functional interaction between the Prp40 WW domains and BBP during splicing.

### AlphaFold modeling places the WW domains in the vicinity of the 5’SS

The lack of a complete Prp40 structure has hindered our understanding of its precise function. To address this, we employed AI-based protein structure prediction using AlphaFold 3 to generate a Prp40 model (Fig. 6A). We then aligned this predicted Prp40 structure within the U1 snRNP in the context of the pre-A complex^29^. Strikingly, this alignment revealed that the Prp40 WW domains, which is followed by an extended unstructured linker, were positioned near the 5’SS and Luc7 (S6). While this proposed positioning of the Prp40 WW domains is consistent with our experimental data, the predicted local distance difference test (pLDDT) score for the unstructured linker is very low (<50) (Fig. 6A). To provide support for the predicted model, we experimentally manipulated the linker length. If the AlphaFold model is correct, shortening the linker should prevent the WW domains reaching the 5’SS, potentially mimicking the observed WW domains-deletion phenotypes. We generated two *prp40* mutants: *prp40-ΔL*, which removes the entire linker region (residues 73-131 deleted) and *prp40-sL*, which shortens the linker region (residues 73-113 deleted) (Fig. 6B, S7). As expected, *prp40-ΔL* resulted in a growth reduction when combined with the *mud2Δ* (Fig. 6C), reminiscent of the *prp40*-*ΔN* phenotype from the SGA genetic screen. This growth reduction was most obvious during the early stage (Day 1) of incubation (Fig. 6C). Interestingly, the *prp40-ΔL* mutant displayed a severe temperature-sensitive phenotype, performing worse than the *prp40*-*ΔN* mutant at 37°C (Fig. 6C; right panel, rows 7 vs. 8 and 9). In contrast, the *prp40*-*sL* mutant has a growth phenotype similar to the wild-type (Fig. 6C; right panel; row 10), suggesting that the 19 amino acids in this linker may be sufficient to properly position the WW domains. We further measured the splicing efficiencies in the *prp40-ΔL* and *prp40-sL* mutants. Indeed, unspliced introns of the 5’SS variants accumulated in the *prp40-ΔL* mutant (Fig. 6D). Yet, splicing of the BS variants was less pronouncedly affected in the *prp40-ΔL* mutant (Fig. 6D), suggesting that a Prp40 devoid of the linker region may still interact with BBP, although less efficiently. Taken together, our data raise a scenario that the functionally critical linker region is structurally flexible enough to allow the WW domains to dynamically interact with the intron 5’SS and BP within the spliceosomal environment.

**Fig. 6.**
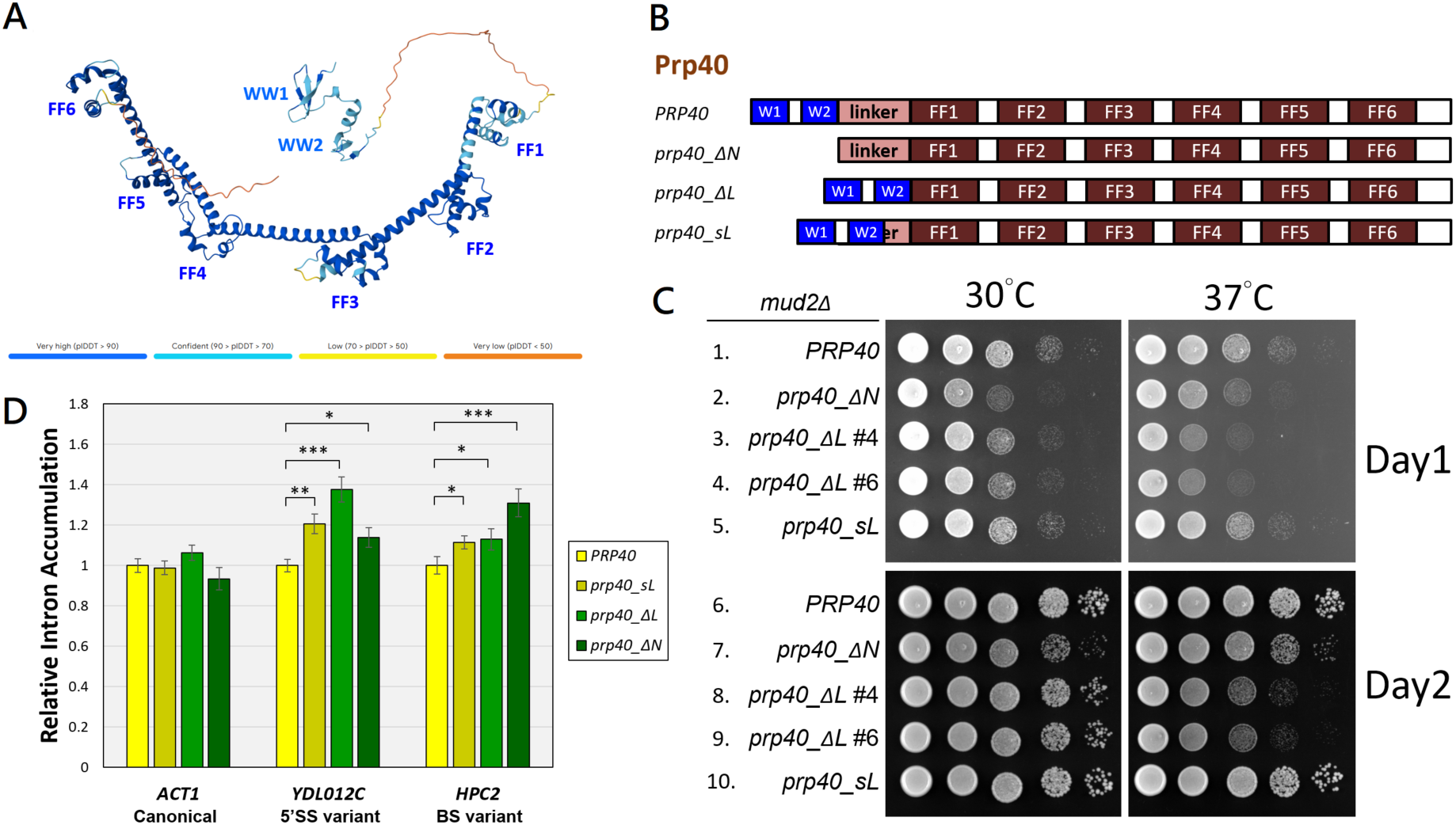
Prp40’s disordered linker may endow WW domains with structural and positional flexibility in relation to 5’SS and BP. (A) AlphaFold 3-predicted Prp40 structure. Prediction confidences are represented by pLDDT scores as shown in color gradient. (B) A schematic diagram of three engineered Prp40 variants. ΔL, linker deletion; sL, a shortened linker in which the two-thirds of the linker is deleted. (C) Growth phenotypes of Prp40 linker-deleted or - truncated strains in the *mud2Δ* genetic background. Our SGA screen uncovered that *mud2Δ* deletion genetically interacts with *WWΔ* (see Results and Table 1). Likewise, complete deletion of the linker (*ΔL*) in the *mud2Δ* genetic background resulted in growth defect at 37°C (rows 3, 4, 8, and 9; right panels), whereas partially truncated linker (*sL*) did not (rows 5 and 10). Cells were serially diluted and spotted onto YPD plates and incubated at the specified temperatures. Images were taken on day 1 and day 2 after incubation. *ΔL* #4 and *ΔL* #6, two independently constructed clones. (D) Prp40 linker-deletion reduces the 5’SS variant intron’s splicing efficiency. Relative intron levels of selected transcripts (canonical, 5’SS, and BS variants) were measured in wild-type (*PRP40*), shortened-linker (sL), linker-deleted (ΔL), and WW domain–deleted (*prp40*-ΔN) yeast strains. N = 5 or 6 biological replicates; error bars are ±SEM; *\*P* < 0.05*, **P* < 0.01, and *\*\*\*P* < 0.001.

## Discussion

The spliceosome faces a fundamental challenge in distinguishing true splice sites from non-informative sequences within pre-mRNA. Despite most of the splice-site sequences in yeast are conserved, the presence of non-canonical 5’SSs and BSs, which represent nearly 40% of the intron-containing genes in the introns, raises the question of how the spliceosome deals with those variations. Our RNA-seq analysis supports the notion that introns with non-canonical splice-site sequences are, in general, spliced less efficiently than canonical ones (Fig 2B). While the sequencing depth of our dataset did not allow us to confidently distinguish splicing efficiency differences between wild-type and WW-domain-deletion backgrounds, the overall trend reinforces our hypothesis that non-canonical introns represent suboptimal substrates for the spliceosome. These observations underscore the need for mechanisms—such as protein-protein interactions mediated by Prp40’s WW domains—to support splicing under conditions of weak *cis*-information. Our study demonstrates that the evolutionarily conserved WW domains of Prp40 play a crucial role in enabling the splicing machinery to accommodate this sequence diversity.

Although previous studies suggested that Prp40 WW domains are dispensable under standard growth conditions^27^, our work revisits this question with a specific focus on intron sequence variation. We show that deletion of both WW domains (*prp40-ΔN*) selectively impairs splicing of introns harboring non-canonical 5′SS or BS sequences, without affecting introns with canonical sequences. The reduced co-transcriptional recruitment of U1 snRNP and BBP to such introns in the *ΔN* mutant background highlights a role for Prp40 in stabilizing early spliceosomal complexes when substrate recognition is suboptimal.

This stabilization may be mediated through physical bridging interactions between the U1 snRNP (via Luc7) and BBP, as previously hypothesized^11,12,31,45^. Our data support this model and further reveal that Prp40’s WW domains are especially important under stress conditions (e.g., low temperature) or when splice-site signals are weak. In this way, the WW domains act as fidelity enhancers, allowing the spliceosome to retain flexibility in substrate recognition without compromising accuracy.

Previous studies showed that, in *Schizosaccharomyces pombe* and mammals, efficient splicing of non-canonical introns requires phosphorylation of branchpoint-binding proteins (Bpb1 in *S. pombe*, SF1 in humans) by SR protein kinases such as Dsk1 and SRPK1/2, respectively^46^, to act within a conserved RXXSP phosphorylation motif. This phosphorylation enhances binding affinity to weak or degenerate branch sites, promoting stable early spliceosomal complex formation^46^. Intriguingly, the PPVY motif in *S. cerevisiae* BBP — which interacts with the Prp40 WW domains — lies just downstream of the conserved RXXSP phosphorylation motif (S8). This spatial arrangement raises the possibility that phosphorylation of BBP, in conjunction with WW domain interaction, could cooperatively regulate its association with non-canonical BS sequences. While *S. cerevisiae* BBP phosphorylation remains underexplored, a similar mechanism could explain how early spliceosome assembly adapts to sequence variability. These parallels reinforce the notion that different organisms may use distinct, yet convergent, strategies to ensure the fidelity and flexibility of early splicing decisions.

Our data also suggest functional distinctions between the two WW domains. Deletion of the first WW domain (WW1), but not both, unexpectedly rescued the cold-sensitive phenotype of BBP mutants (*msl*5-S194P and *msl*5-V195D). This rescue was associated with altered BBP recruitment dynamics, implying that the WW1 domain may restrict BBP release from the commitment complex. Conversely, WW2 appears to be required for maintaining BBP stability on non-canonical BS sequences. Thus, the two WW domains may differentially regulate BBP binding and release, adding a layer of spatiotemporal regulation to spliceosome assembly. This nuanced role highlights splicing machinery’s adaptability and suggests that Prp40 fine-tunes the complex dynamics in the face of different splicing substrates.

Yeast introns are under strong selection pressure to maintain splicing-compatible sequences. Canonical sites are preferentially recognized due to high-affinity interactions with U1 snRNP and BBP. However, the existence and efficient processing of non-canonical introns suggest that auxiliary mechanisms exist to ensure their inclusion. Our results support a model in which the Prp40 WW domains serve as one of such auxiliary mechanisms—providing structural and functional support when cis-elements are suboptimal.

Interestingly, in higher eukaryotes, intron diversity is even more pronounced. Although the *S. cerevisiae* Prp40 protein is considered a fungal-specific U1 snRNP protein, its mammalian counterparts (e.g., PRPF40A/B) and other WW-domain-containing splicing regulators such as CA150 interact with SF1 (the human BBP ortholog) and U2AF^16,47^.

These interactions may perform a similar bridging or stabilization function during early spliceosome assembly on degenerate introns. Moreover, mutations in several of these early splice-site recognition proteins—including PRPF40B, SF1, and U2AF65—are implicated in myelodysplastic syndromes and other spliceopathies, underscoring their physiological relevance^26^. Thus, our findings suggest a conserved mechanistic logic: WW-domain-containing proteins may serve as dynamic scaffolds that stabilize early spliceosome complexes, particularly when substrate recognition is weak or degenerate.

Cryo-EM studies of the yeast spliceosome have not revealed the WW domains of Prp40, likely due to structural flexibility^28,29,37^. To overcome this limitation, we employed AlphaFold to model the full-length Prp40 protein. The predicted model revealed a topologically conserved unstructured linker (S1A) connecting the WW and FF domains, allowing the WW domains to potentially reach the 5’SS and interact with U1 snRNP components such as Luc7. Experimental validation of this model, through targeted deletions and linker shortening, confirmed that the linker is required for WW-domain positioning and non-canonical intron splicing. Complete linker deletion impaired splicing of 5′SS variant but less so for BS variant (Fig. 6D), indicating that the linker primarily mediates WW-domain access to U1-bound splice sites, while BBP interaction remains largely intact. This supports a spatially defined and structurally tunable role of the WW domains in spliceosome assembly.

In conclusion, our study redefines the functional importance of the Prp40 WW domains in *S. cerevisiae* by revealing their critical role in facilitating splicing of non-canonical introns. These domains act as structural adaptors that promote stable early complex formation when canonical RNA-RNA interactions are weakened. In the broader evolutionary context, such mechanisms may have evolved to allow eukaryotic genomes to expand their splicing repertoire without compromising fidelity. Going forward, it will be valuable to examine whether other accessory splicing factors similarly enhance non-canonical intron splicing and whether WW-domain-containing proteins in higher eukaryotes serve similar roles. The potential interplay between phosphorylation of BBP (or SF1) and WW-domain interaction also warrants further mechanistic dissection, particularly in metazoans where splicing regulation is highly dynamic and context dependent.

## Materials and Methods

### Yeast strain construction

The heterozygous diploid strain *prp40::KanMX/PRP40* (YTC1125; *MAT***a***/α his3Δ1/his3Δ1 leu2Δ0/leu2Δ0 LYS2/lys2Δ0 met15Δ0/MET15 ura3Δ0/ura3Δ0 prp40::KanMX/PRP40*) was obtained from the Yeast Knockout Collection (Open Biosystems, now Horizon Discovery). To prepare a parental strain suitable for plasmid shuffling, we first introduced a low-copy *PRP40* plasmid carrying *PRP40/URA3/CEN* (pPRP40-2) into YTC1125. The resulting diploid was then sporulated and tetrads were dissected. From these, we selected an α-haploid isolate (YTC1125_32) that carries the chromosomal *prp40::KanMX* allele and is complemented by the plasmid pPRP40-2. This isolate was subsequently used as the parental strain for plasmid shuffling.

To generate the *PRP40*, *prp40-ΔWW1*, *prp40-ΔWW2*, and *prp40-ΔN* strains, the low-copy plasmids pPRP40-1 (*PRP40/LEU2/CEN*), pPRP40-32 (*prp40-ΔWW1/LEU2/CEN*), pPRP40-33 (*prp40-ΔWW2/LEU2/CEN*), and pPRP40-34 (*prp40-ΔN/LEU2/CEN*) were introduced to replace the complementing pPRP40-2. This yielded the strains YTC1125_69 (*PRP40*), YTC1125_70 (*ΔWW1*), YTC1125_71 (*ΔWW2*), and YTC1125_72 (*ΔN*). Similarly, the *prp40_sL* (YTC1125_512) and *prp40-ΔL* (YTC1125_502) strains were constructed by replacing pPRP40-2 in YTC1125_32 with pPRP40-36 (*prp40_sL/LEU2/CEN*) and pPRP40-37 (*prp40-ΔL/LEU2/CEN*), respectively.

To generate these same *PRP40* and mutant alleles in the BBP variant background, YTC1125_32 was crossed with YTC1122 (*MAT***a** *his3Δ1 leu2Δ0 met15Δ0 ura3Δ0 msl5-S194P-TAP-HIS3*) and YTC1196 (*MAT***a** *his3Δ1 leu2Δ0 met15Δ0 ura3Δ0 msl5-V195D-TAP-HIS3*). Diploids were sporulated, and tetrad were dissected to obtain haploid strains (*prp40::KanMX msl5-S194P-TAP-HIS3* and *prp40::KanMX msl5-V195D-TAP-HIS3*) that carried the pPRP40-2 plasmid. By plasmid shuffling, pPRP40-1, pPRP40-32, pPRP40-33, and pPRP40-34 were introduced to replace pPRP40-2, generating the corresponding allelic series in the BBP variant background.

To generate strains for ChIP analysis, YTC1125_32 was crossed with YTC1148 (*MAT***a** *his3Δ1 leu2Δ0 met15Δ0 ura3Δ0 YHC1-TAP-HIS3)* and YTC1030 (*MAT***a** *his3Δ1 leu2Δ0 met15Δ0 ura3Δ0 MSL5-TAP-HIS3*). Diploids were sporulated, and tetrad were dissected to obtain the haploid strains YTC1148_148 (*prp40::KanMX YHC1-TAP-HIS3 PRP40/URA3/CEN*) and YTC1030_126 (*prp40::KanMX MSL5-TAP-HIS3 PRP40/URA3/CEN*). After plasmid shuffling, pPRP40-1 and pPRP40-34 were introduced to replace pPRP40-2, generating *PRP40* and *prp40-ΔN* derivatives. Thus, the final strains used in the ChIP experiments were YTC1148_151 (*YHC1-TAP-HIS3 PRP40*) and YTC1148_154 (*YHC1-TAP-HIS3 prp40-ΔN*)] in the *YHC1-TAP* background, and YTC1030_133 (*MSL5-TAP - HIS3 PRP40*) and YTC1030_136 (*MSL5-TAP - HIS3 prp40-ΔN*) in the *MSL5-TAP* background.

### Competitive fitness assay

To detect the relative fitness of the query strains, overnight cultures of the query strains and the reference GFP-expressing strain (YTC1313) were refreshed by 1:60 dilution in YPD (1% yeast extract, 2% peptone, and 2% glucose) medium at 30°C for 3 hr. Next, 5 x 10^6^ cells of early-log-phase cultures of query strains were mixed with equal number of the GFP reference cells in 1 ml of sterile H_2_O. From this mixture, about 10^4^ mixed cells were transferred into 3 ml of YPD for another 12-hr growth at 30°C, allowing approximately six generation of competitive growth. In addition, 1000 cells from this mixture (1:1 query vs. reference) were immediately analyzed by a fluorescence-activated cell sorter (FACS Calibur^TM^, Becton Dickinson, Franklin Lakes, NJ), the reading of which was set as the Time 0 data. The number of GFP cells in the samples (1000 cells) at Time 12 hr was also analyzed by the same method.

### Synthetic genetic array (SGA)

To generate a parental strain for SGA screening, the *prp40::KanMX* allele in YTC1125_32 was replaced with *prp40::clonNAT* by homologous recombination using a PCR-amplified *clonNAT* fragment, yielding strain YTC1125_219. The low-copy plasmids pPRP40-1 (*PRP40/LEU2/CEN*), pPRP40-32 (*prp40-ΔWW1/LEU2/CEN*), pPRP40-33 (*prp40-ΔWW2/LEU2/CEN*), and pPRP40-34 (*prp40-ΔN/LEU2/CEN*) were then used to replace pPRP40-2 in YTC1125_219, generating the SGA query strains YTC1125_225 (*PRP40*), YTC1125_226 (*ΔWW1*), YTC1125_227 (*ΔWW2*), and YTC1125_228 (*ΔN*).

To identify synthetic genetic interactions between *PRP40* alleles and the collection of nonessential gene deletions, we followed established SGA methodology^34^ with minor modification. Briefly, the “magic marker” (*can1Δ::STE2pr-Sp_his5 lyp1Δ::STE3pr-LEU2*) from strain YTC1686 (Y8205; a gift from C. Boone) was introduced into ∼4,600 *MAT***a** nonessential deletion strains (the “Standard SGA Array”, also a gift from C. Boone) through mating, diploid selection, and haploid isolation^34^. This modification enabled selective recovery of the desired haploid mating type in subsequent crosses with the *PRP40* query strains. With modest adjustments, the SGA protocol was carried out using automated pinning using a RoToR HDA robot (Singer Instruments)^33,34^. Each of the four query strains was individually crossed with the magic-marker-modified deletion collection on YPD plates overnight. Diploids were selected twice on YPD plates supplemented with G418 (200 mg/L; Sigma A1720) and clonNAT (100 mg/L; Jena Bioscience AB-102), then sporulated at 22°C for 5 days. *MAT***a** haploids carrying both *prp40::clonNAT* and the respective nonessential deletions were subsequently recovered following SGA procedures. Colony growth images were captured and analyzed using ScreenMill^48^.

### Splicing-sensitive microarray analysis

Sample collection and splicing microarray analysis were performed exactly as described^40,49^. Briefly, strains YTC1125_69 (*PRP40*) and YTC1125_72 (*prp40-ΔN*) were grown in YPD at 30°C to an optical density (OD) of A_600_ = 0.7. Total cellular RNA was extracted and reverse transcribed into Cy3- and Cy5-labeled cDNAs by random priming. These cDNA samples were hybridized to splicing microarrays, yielding raw data that were processed into three datasets: P (pre-mRNA), M (mRNA), and T (total transcripts), representing the differential hybridization signals of 249 intron-containing genes. For each array, dye-swap replicates were included. The splicing microarray data have been deposited in GEO under accession number **GSExxxx** (Note: Because of current government shutdown, NCBI/GEO are not accepting or processing any submissions. We will update the GEO information when it is resolved).

### RNA-seq analysis

*PRP40* yeast cells (YTC1125_69) were cultured in 50 ml of liquid YPD medium at 30°C to mid-log phase (OD_600_ ≈ 0.8). RNA was extracted using MasterPure^TM^ Yeast RNA Purification Kit (Lucigen/Epicentre now Cambio, MPY03100) and subsequently treated with the Ribo-Zero^TM^ Magnetic Gold Kit (Yeast) (Epicentre, now illumine, MRZY1324) to remove ribosomal RNA following manufacturer’s instructions. Libraries were prepared using the TruSeq Stranded mRNA Preparation Kit (Illumina), and deep sequencing was performed on an Illumina HiSeq2500 at the NGS High-Throughput Genomics Core, Biodiversity Research Center, Academia Sinica. The sequencing data has been deposited in GEO under accession number GSE308638.

### *In vivo* splicing analyses

YTC1125_69 (*PRP40*) and YTC1125_72 (*prp40-ΔN*) cells were cultured in 50 ml of YPD at 30°C to mid-log phase (OD_600_ ≈ 0.7). Total RNA was extracted using the MasterPure^TM^ Yeast RNA Purification Kit (Lucigen, now Biosearch Technologies; MPY03100) and reverse-transcribed to cDNA with the Maxima First Strand cDNA Synthesis Kit for RT-qPCR (Thermo Scientific^TM^ K1641). Splicing efficiency of a selected intron-contain gene was assessed by qPCR using two sets of gene-specific primers to quantify unspliced (pre-mRNA, P) and spliced (matured mRNA, M) RNA species. Quantitative PCR was performed on an Applied Biosystems StepOnePlus^TM^ Real-Time PCR System using Fast SYBR™ Green Master Mix (Applied Biosystems, 4385612). To minimize pipetting errors, reactions were assembled using an automated pipetting system (EzMate 401, Arise Biotech). The unspliced species were detected with primers In-Forward/Ex-Reverse spanning the intron-exon 2 boundary, while the spliced species were detected with primers ExJ-Forward/Ex-Reverse spanning the exon-exon junction and exon 2 (S5 Table). The intron accumulation index was calculated as the ratio of unspliced (P) over spliced species (M), normalized to the corresponding index obtained from the wild-type strain. Each experiment was performed in triplicate and error bars represent the standard error of the mean (SEM).

### Splicing reporter analysis

The splicing reporter plasmid (*ACT1-CUP1*/p424GPD) was a gift from Jon Staley^50^. Splice-site variants derived from this plasmid were generated by site-directed mutagenesis PCR using a high-fidelity DNA polymerase (Everything Biotech E-Bio Fusion). To construct the testing strain, the chromosomal *PRP40* allele in strain YTC639 (*MATα cup1::ura3 ura3 leu2 trp1 lys2 ade2 his3 GAL+*) (yJPS11, a gift from Jon Staley) was disrupted using a PCR-amplified *prp40::KanMX* fragment and complemented with pPRP40-2 (*PRP40/URA3/CEN*). This plasmid was subsequently replaced with pPRP40-1 (*PRP40/LEU2/CEN*), pPRP40-32 (*prp40-ΔWW1/LEU2/CEN*), pPRP40-33 (*prp40-ΔWW2/LEU2/CEN*), or pPRP40-34 (*prp40-ΔN/LEU2/CEN*), yielding strains YTC639_297(*PRP40*), YTC639_305 (*prp40-ΔWW1*), YTC639_306 (*prp40-ΔWW2*), and YTC639_298 (*prp40-ΔN*), respectively. Following introduction of reporter plasmids carrying 5′SS or BS variants, cells were spotted onto selective synthetic complete plates (SC-Trp) supplemented with varying concentrations of CuSO_4_, as described in^44^, and incubated at various temperatures.

### Chromatin IP and real-time PCR

We followed an established ChIP procedure^51^ to monitor co-transcriptional recruitment of U1 snRNP and BBP to nascent transcripts. Two yeast strains (see above) were used for this purpose, in which U1C and BBP were individually was tagged with TAP. Briefly, 50 ml of yeast cells at OD_600_ ≈ 0.8 were cross-linked with 1% formaldehyde (Sigma Aldrich F8775) at room temperature for 10 min. Cells were then resuspended in ChIP buffer (50 mM HEPES pH 7.5, 140 mM NaCl, 1% Triton X-100, 0.1% DOC, 0.1% SDS, and 1X protease inhibitor; Roche cOmplete 11836145001) with 250 µl of glass beads, and disrupted using a Mini-Beadbeater-16 (BioSpec) for 2 min. The insoluble cross-linked fraction was resuspended in ChIP buffer and sonicated with a Q800R2 Sonicator System (Qsonica) to shear DNA to ∼300 bp fragments. Immunoprecipitation of U1C-TAP or BBP-TAP was carried out with IgG Sepharose 6 Fast Flow affinity resin (Cytiva 17096901) at 4°C overnight. After standard wash procedures, the crosslinks were reversed and DNA extracted. Primer pairs spanning the entire query genes were used for qPCR on an Applied Biosystems StepOnePlus Real-Time PCR System with Fast SYBR™ Green Master Mix (Applied Biosystems, 4385612). Data represent the mean of three independent experiments, with standard error of the mean (SEM) indicated. Enrichment of the U1 snRNP and BBP was normalized to the first primer pair.

### Quantification and statistical analysis

Statistical analysis was conducted using Microsoft Excel. Statistical significance was assessed using an unpaired, two-tailed Student’s t test. P-value summary for all statistical tests used in this study is: ns, P-value > 0.05; ∗ P-value < 0.05; ∗∗ P value < 0.01; ∗∗∗ P-value < 0.001.

### Cryo-EM density map retrieval and analysis

The predicted atomic models of Prp40 and Luc7 were generated using the AlphaFold 3 server^30^. The cryo-EM density map of the spliceosomal pre-A complex (EMD-13033) and its associated atomic model (PDB ID: 7OQE) were retrieved from the Electron Microscopy Data Bank (https://www.ebi.ac.uk/pdbe/emdb/) and Protein Data Bank (https://www.rcsb.org), respectively. All models and the cryo-EM map were visualized using UCSF ChimeraX^52^. The AlphaFold predicted Prp40 model was docked into the cryo-EM density map using the ChimeraX “Fit in Map” tool. Rigid-body fitting was performed to align the predicted helical regions with the corresponding density features to achieve maximum coverage, leading to the re-assignment of consecutive helices (FF1–FF4) based on the fit. The contour threshold for the cryo-EM map was initially set to 0.0140 to show a tight coverage with the deposited model. This threshold was then systematically relaxed to 0.008 to reveal additional masked density, which progressively enveloped the FF6 region of the predicted Prp40 model.

## Acknowledgments

We thank P. Siliciano for providing anti-Prp40 antibody, J. P. Staley for the yeast strain and the plasmid used in the splicing reporter assay, and J.-Y. Leu for the GFP-expression reference strain. We thank members of the Chang laboratory for critical discussions. T.-H. C. was supported by Ministry of Science and Technology (MOST 110-2311-B-001-042) and by Academia Sinica Grand Challenge Seed Grant (AS-GCS-110-03) and Innovative Research Project (AS-IR-112-02-A).

## Author contributions

T.-H.C. and L.T. conceived and led the project, designed the experiments, analyzed the data, and wrote the manuscript. L.T., C.-S.Y., S.-L.C. and F.-L.Y. performed the experiments. H.-K.W., H.-Y.T., and L.T. analyzed the RNA-seq data. A.L. and J.A.P. performed the microarray experiments and analyzed the corresponding data. H.-H.L., L.T., and W.-H.C. conducted the structural analysis. C.-L.H. and C.-F.K. provided reagents.

## Declaration of generative AI and AI-assisted technologies in the manuscript preparation process

During the preparation of this work we used AlphaFold 3 to predict protein structures. After using this tool, we reviewed and edited the content as needed and take full responsibility for the content of the published article.

**Figure S1.**
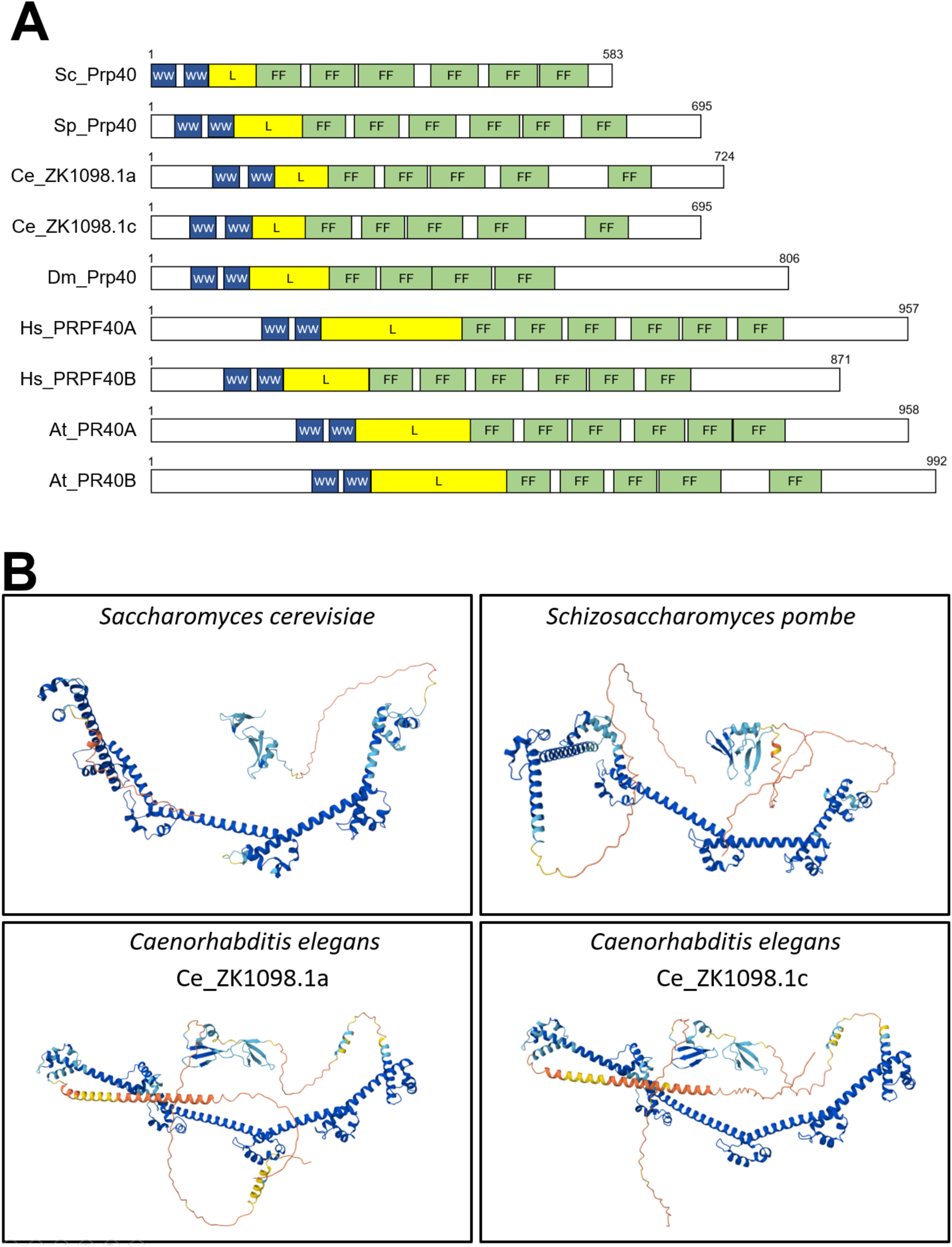
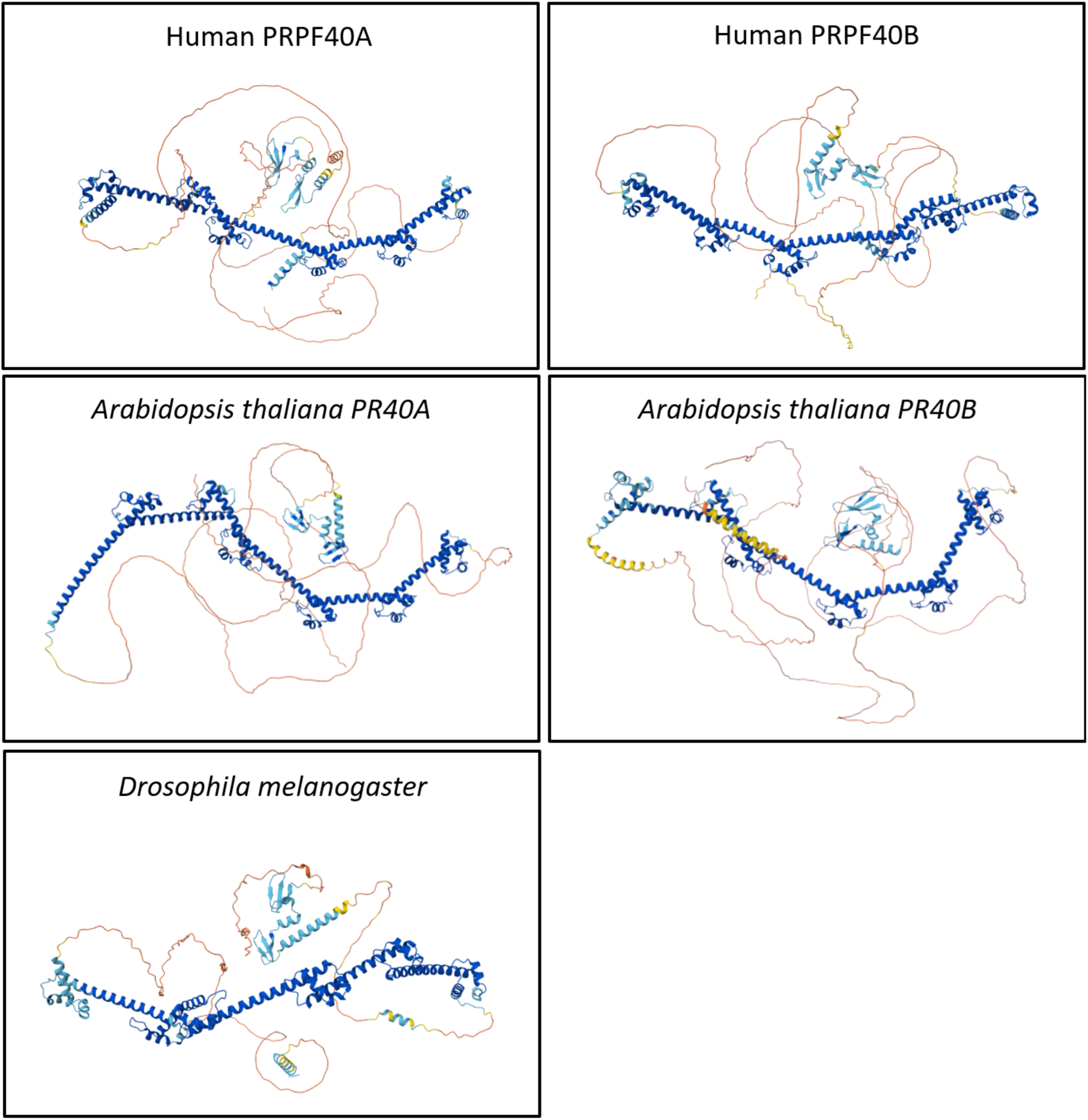
Conserved domain architecture alignment and AlphaFold 3-predicted structures of Prp40 homologs across diverse eukaryotes. **(A)** Conserved domain architecture of Prp40 homologs across eukaryotes. Schematic representation of the conserved domain organization of Prp40 orthologs: *Saccharomyces cerevisiae* (Sc_Prp40), *Schizosaccharomyces pombe* (Sp_Prp40), *Caenorhabditis elegans* (Ce_ZK1098.1a and Ce_ZK1098.1c), *Drosophila melanogaster* (Dm_Prp40), *Homo sapiens* (Hs_PRPF40A and HS_PRPF40B), and *Arabidopsis thaliana* (At_PR40A and At_PR40B). The alignment highlights the conservation of two WW domains at the N-terminus and multiple FF domains throughout the protein. An unstructured linker region (yellow box; L), located between the WW and FF domains, is also topologically conserved across species, suggesting a potential regulatory or scaffolding role. Despite variations in protein length and domain number, the overall architecture is preserved, indicating the functional importance of these domains in Prp40-mediated RNA splicing processes. The total amino acid length of each homolog is labeled in the figure. **(B)** AlphaFold 3-predicted structures of evolutionarily conserved Prp40 orthologs across diverse eukaryotes. Predicted three-dimensional structures of Prp40 orthologs generated using AlphaFold 3 are shown for *Saccharomyces cerevisiae* (Sc_Prp40), *Schizosaccharomyces pombe* (Sp_Prp40), *Caenorhabditis elegans* (Ce_ZK1098.1a and Ce_ZK1098.1c), *Drosophila melanogaster* (Dm_Prp40), *Homo sapiens* (Hs_PRPF40A and HS_PRPF40B), and *Arabidopsis thaliana* (At_PR40A and At_PR40B). The structures illustrate a conserved overall organization characterized by N-terminal WW domains and multiple FF domains arranged along an extended scaffold. Notably, an intrinsically disordered region is predicted between the WW and FF domains in all orthologs, suggesting a conserved functional flexibility or interaction platform. Despite species-specific variations in length and domain spacing, the core structural elements are maintained, reflecting evolutionary pressure to preserve Prp40’s role in spliceosome assembly and pre-mRNA processing.

**Figure S2.**
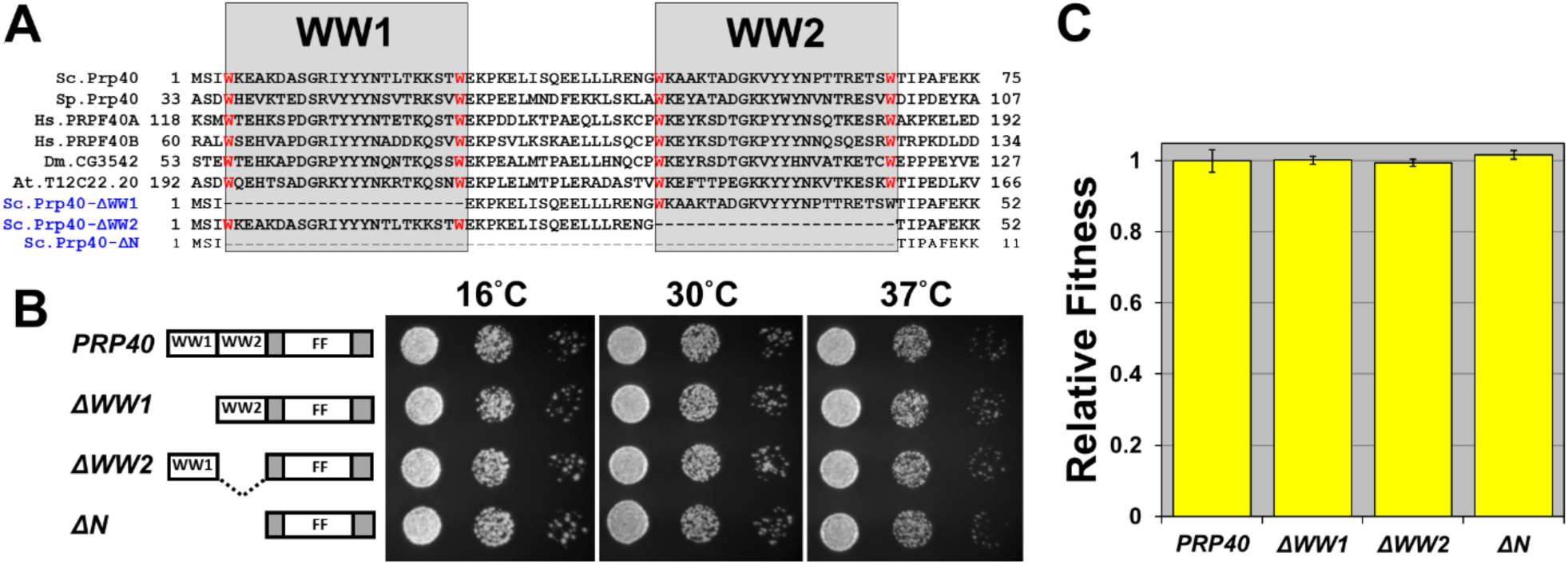
Growth phenotypes upon deletion of Prp40 WW domains. **(A)** Schematic representation of the strategy used to delete the WW domain(s) of *Saccharomyces cerevisiae* Prp40. **(B)** Spotting assay of yeast strains carrying different WW domain deletions. Cultures were subjected to 10-fold serial dilutions, and equal volumes were spotted onto YPD plates and incubated at the indicated temperatures. **(C)** Competitive fitness assay of the indicated *PRP40* WW-domain deletion strains in YPD.

**Figure S3.**
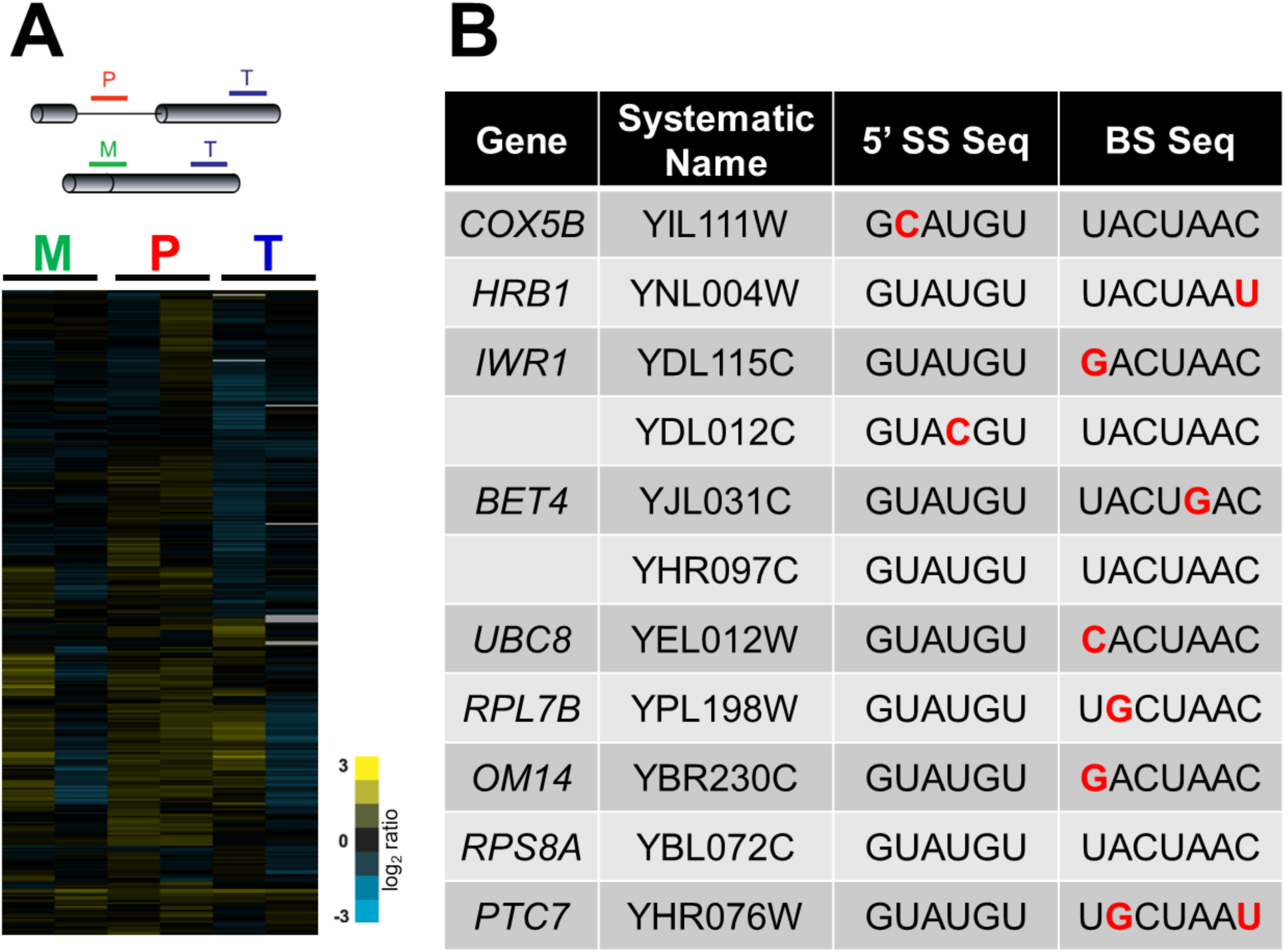
Splicing-specific microarray assay. RNA was extracted from wild-type and *prp40* WW-domain deletion yeast strains and analyzed using a splicing-specific microarray (developed by Pleiss^1–3)^. Primers targeting intronic sequences (P, precursor), exon–exon junctions (M, mature), and exon 2 (T, total) are indicated in panel **(A)**. The lower portion of panel (A) shows results from two biological replicates, representing the RNA signal intensity detected by each oligonucleotide probe. RNA species showing intron enrichment are listed in panel **(B)**, ordered from highest to lowest abundance. Non-canonical 5′SS and BS sequences are highlighted in red.

**Figure S4.**
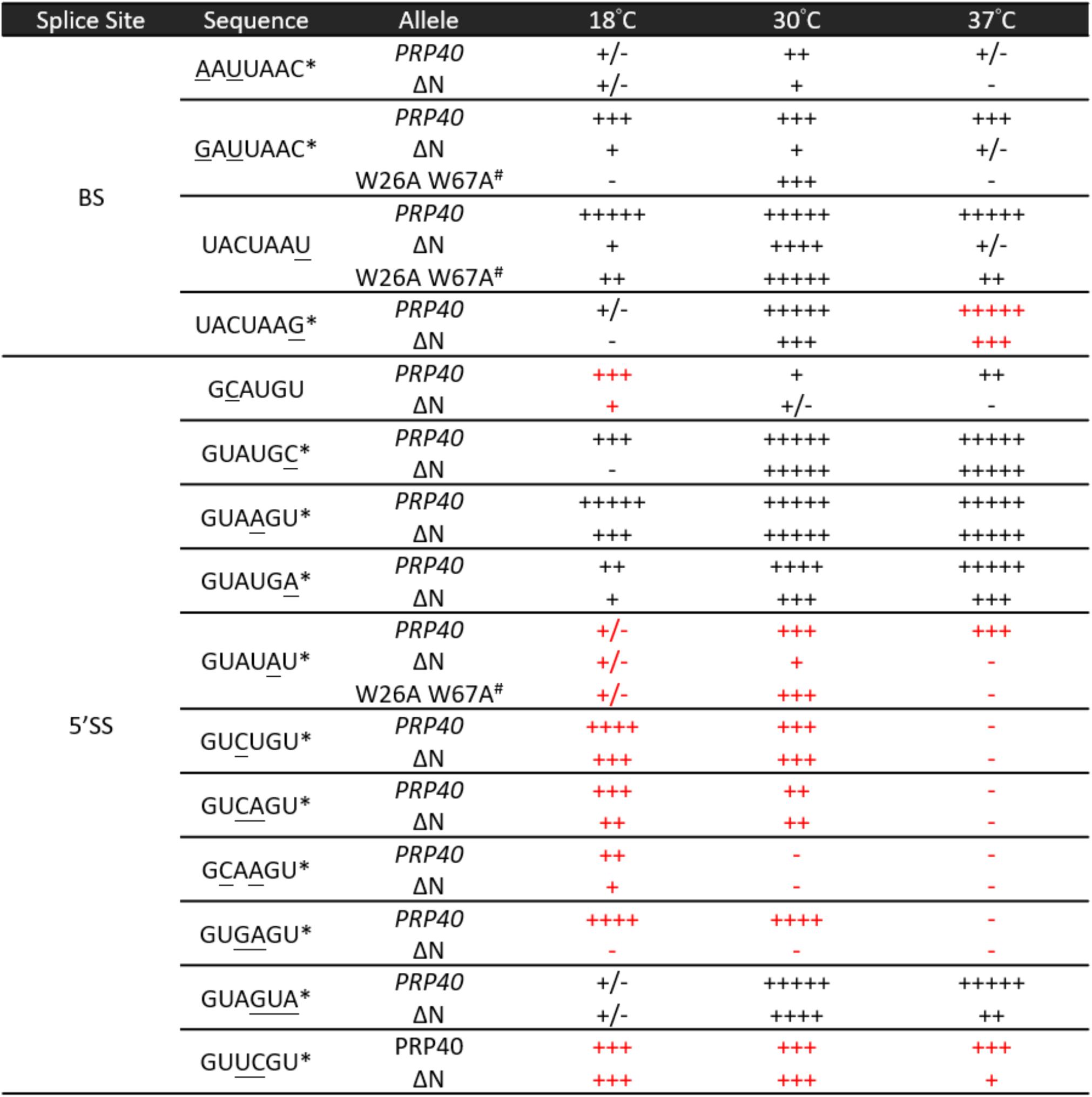
*In vivo* splicing screen result. · No difference BS: AACUAAC, GACUAAC, CACUAAC, UGCUAAC, UAUUAAC, UACUGAC, UAGCAAC, UGCUAAU, UGGUAAG; 5’SS: GUACGU, GUUAAG · Lethal BS: UACUACC; 5’SS: AUUUGU, GUAUGG, GUUUGU · Red: tested on 0.1 mM of CuSO4; Black: tested on 0.5 mM of CuSO4 · *: 13 new WW-dependent non-canonical BSs and 5’SSs identified in this screen · #: The *prp40*-W26A W67A allele has been engineered so that the conserved tryptophan residues in the WW domain are replaced with alanines.

**Figure S5.**
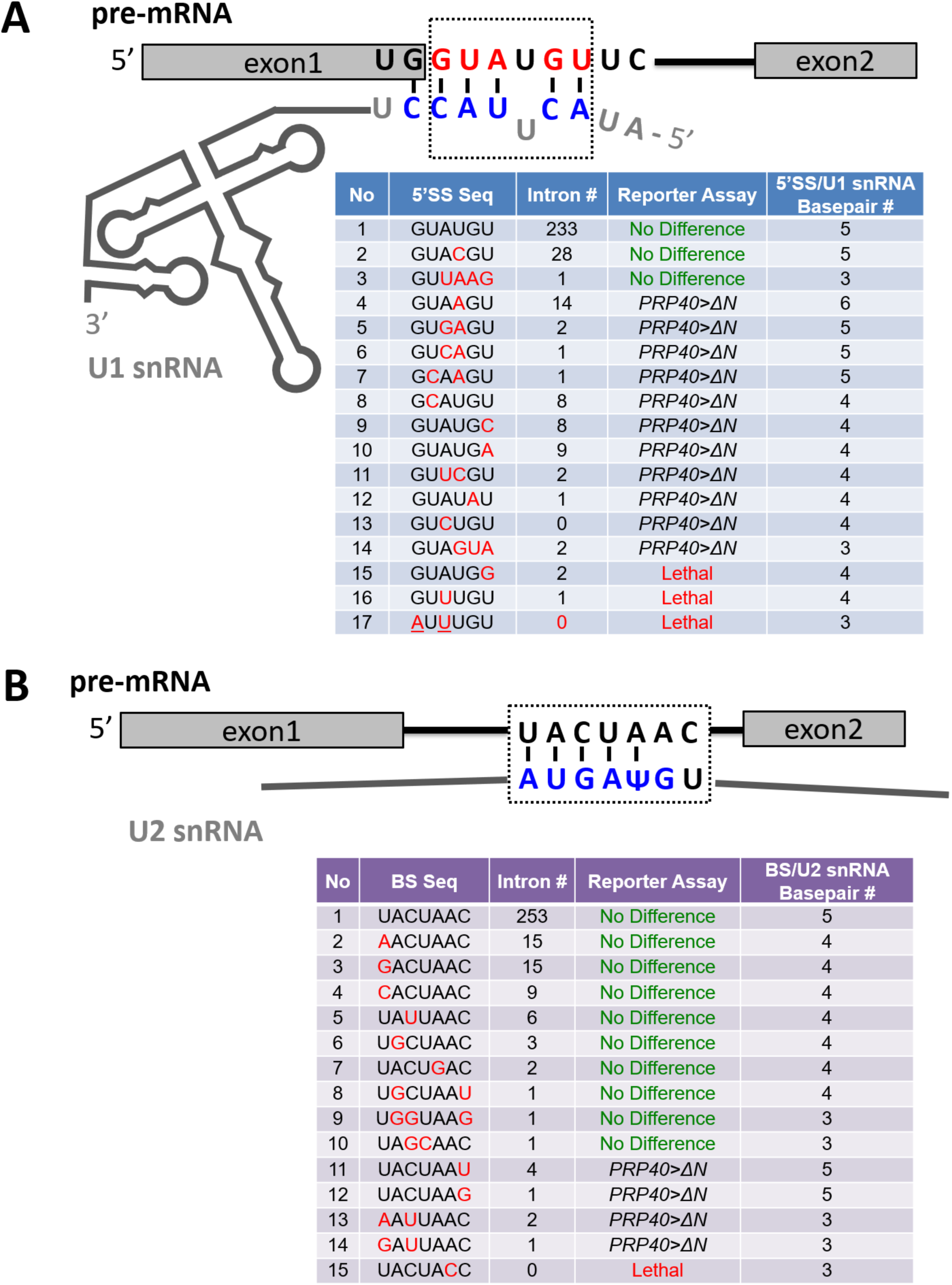
Comparison of non-canonical 5′SS and BS base pairing with U1 and U2 snRNAs. **(A)** Comparison of non-canonical 5′SS variants tested in the reporter assay, showing the extent of base pairing between each 5′SS and U1 snRNA. **(B)** Comparison of non-canonical BS variants tested in the reporter assay and their base-pairing interactions with U2 snRNA. “*PRP40>*Δ*N*” indicates that yeast exhibit enhanced growth in the wild-type *PRP40* background relative to the *ΔN* background.

**Figure S6.**
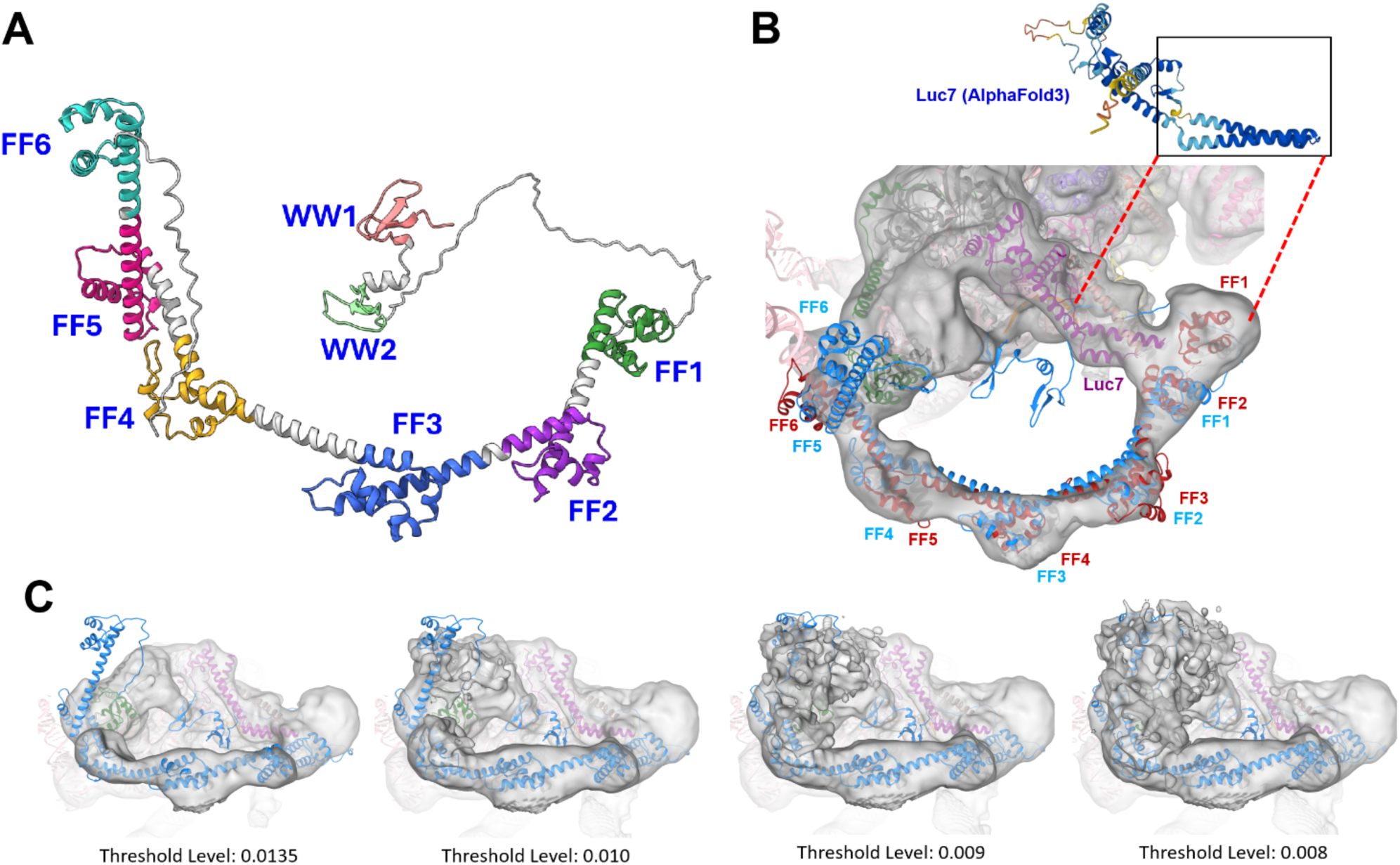
Structure-based interpretation for potential communications between Prp40 WW domains and U1 snRNP core. **(A)** A complete atomic model of Prp40 was predicted from its sequence using AlphaFold 3^4^, and its FF domains were manually colored in ChimeraX. The major features of this model contain five long α-helices (FF1-FF5) and one short α-helix (FF6). It is noted that the sequences that connect WW2 and FF1 is predicted to be disordered, which renders the positioning of the WW domains indefinite. However, it does allow WW domains to physically communicate with Luc7 in the core. **(B)** The original cryo-EM map from Lührmann’s group (EMD-13033)^5^, contoured at a threshold of 0.0135, assigned all six FF domains (colored in red). However, AlphaFold3 predictions indicate that the region previously identified as FF1 is in fact Luc7 (see comparison with the AlphaFold3-predicted Luc7 structure in the inset). Accordingly, the regions designated as FF2-FF6 (red) correspond instead to FF1-FF5 (Blue). The previously unassigned FF6 domain becomes clearly detectable when the cryo-EM map is displayed at a lower threshold of 0.010. Further details are provided in **(C)**. **(C)** A panel that displays different views of the cryo-EM map in **(B)** with various thresholds. When the threshold is set at 0.010, the FF6 helix can be assigned to an upright protrusion that is entirely within the original 3D density map (EMD-13033).

**Figure S7.**
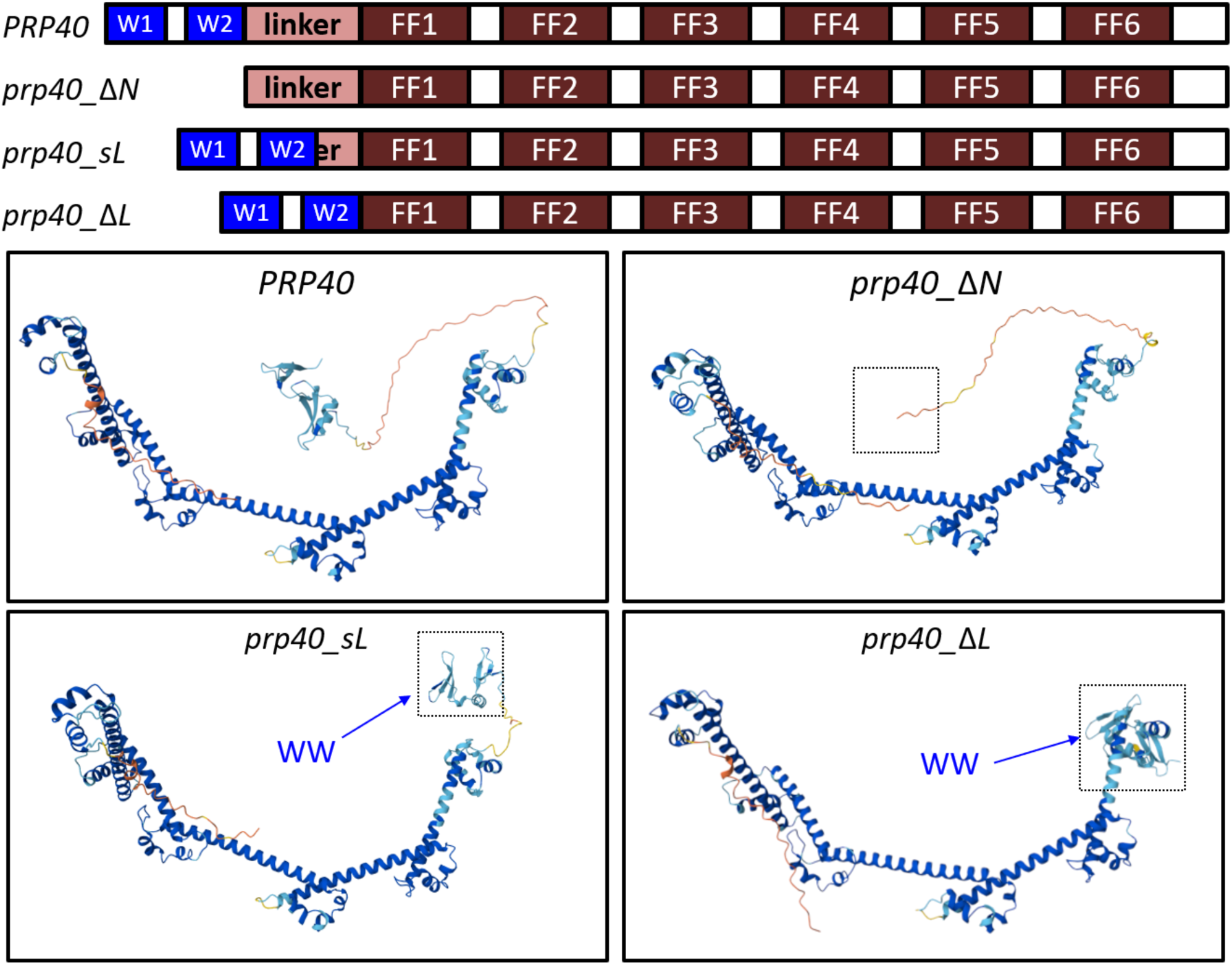
**AlphaFold 3 predicted structures of Prp40 variants.**

**Figure S8.**
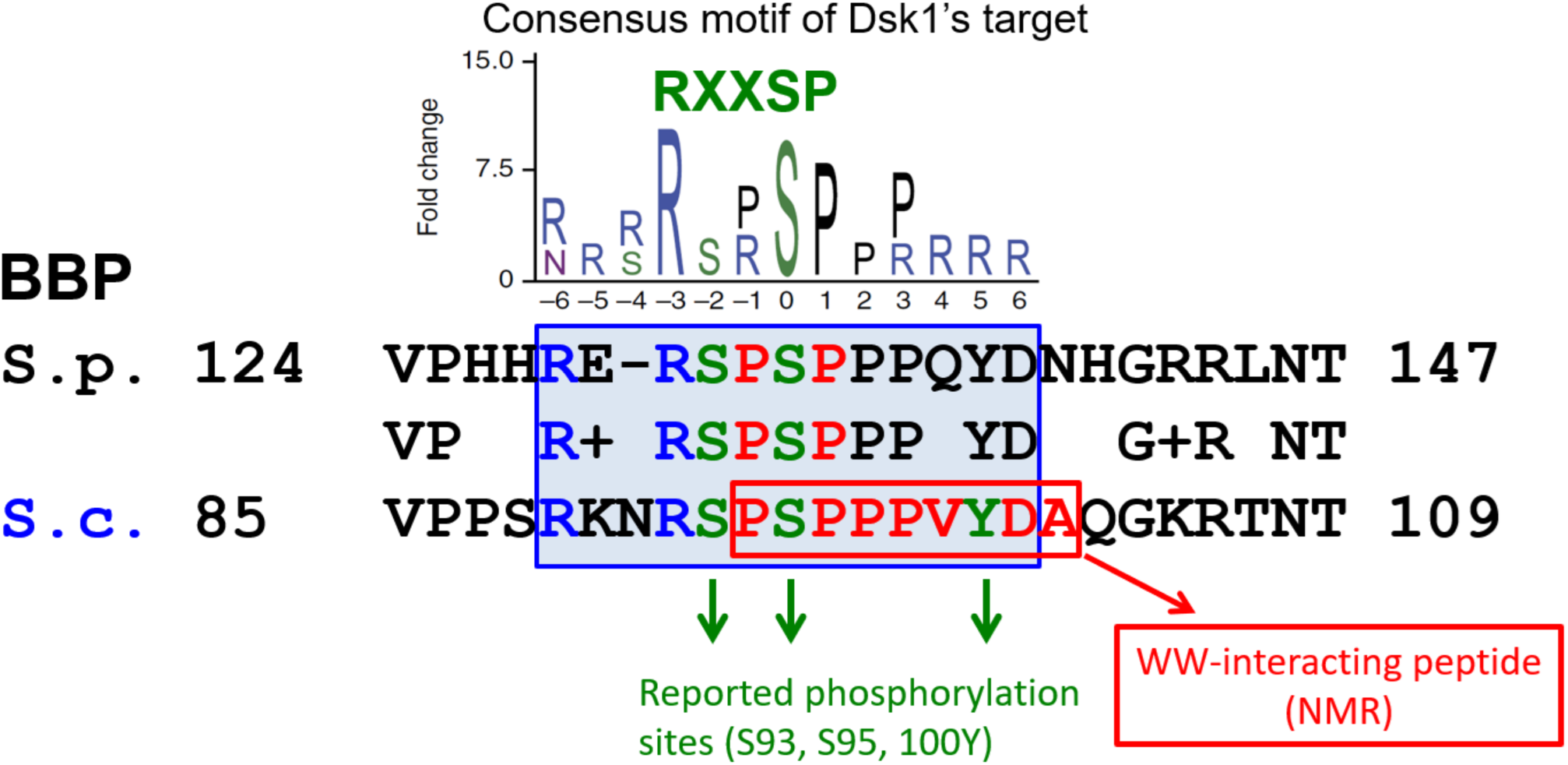
Conservation and Functional Context of the Dsk1 Recognition Motif in Branchpoint-Binding Proteins. The Dsk1 recognition motif is conserved in *S. cerevisiae* and overlaps with the region that interacts with the Prp40 WW domain. The conserved RXXSP phosphorylation motif, recognized by SR protein kinases such as Dsk1, lies immediately upstream of the PPVY sequence that mediates WW-domain binding. The red boxed sequence indicates the peptide used in the NMR structural study^6^ of the Prp40 WW domain. The top chart (Consensus motif of Dsk1’s target) is adapted from Lipp et al.^7^.

---

**Table S1:**
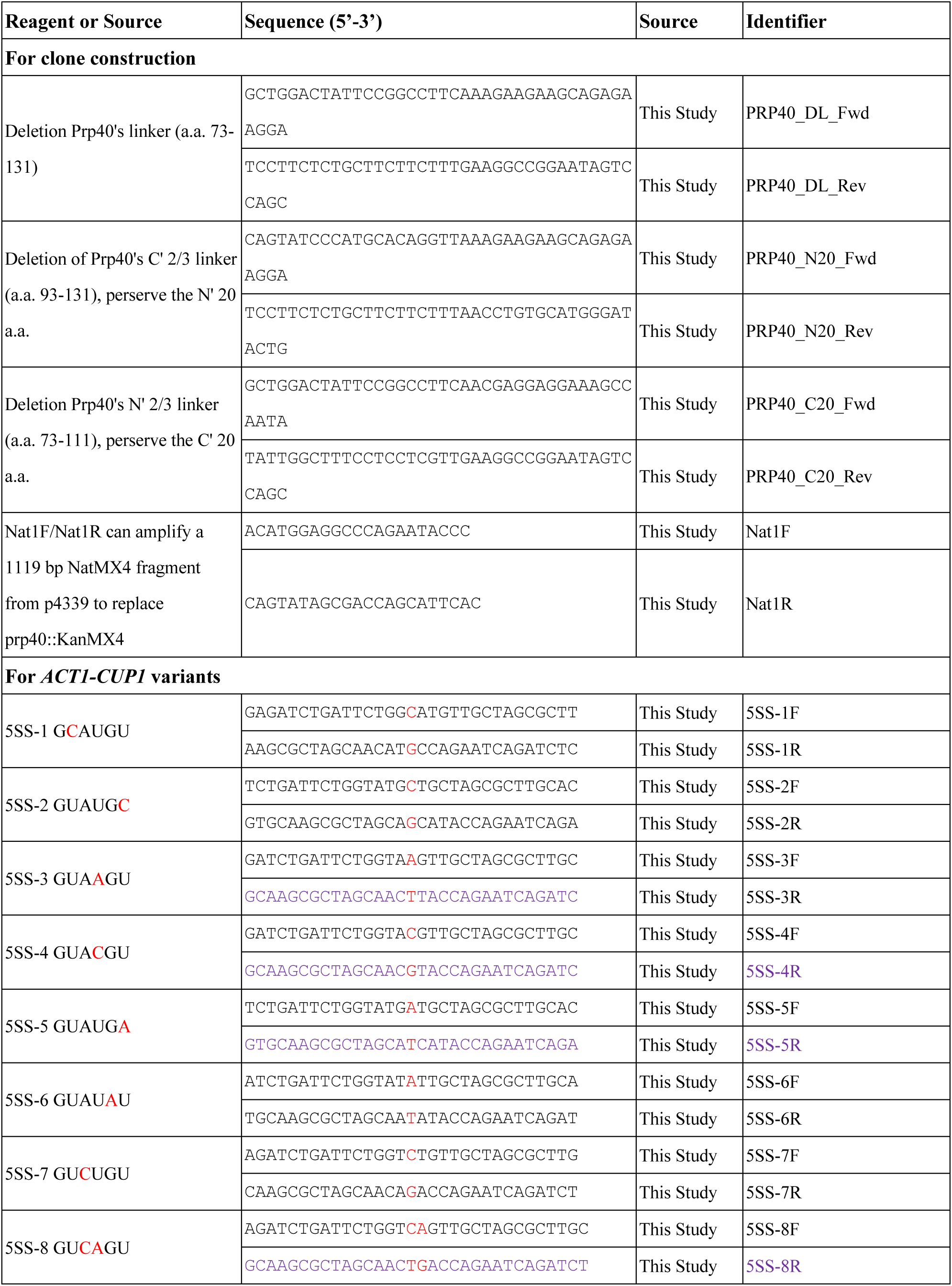
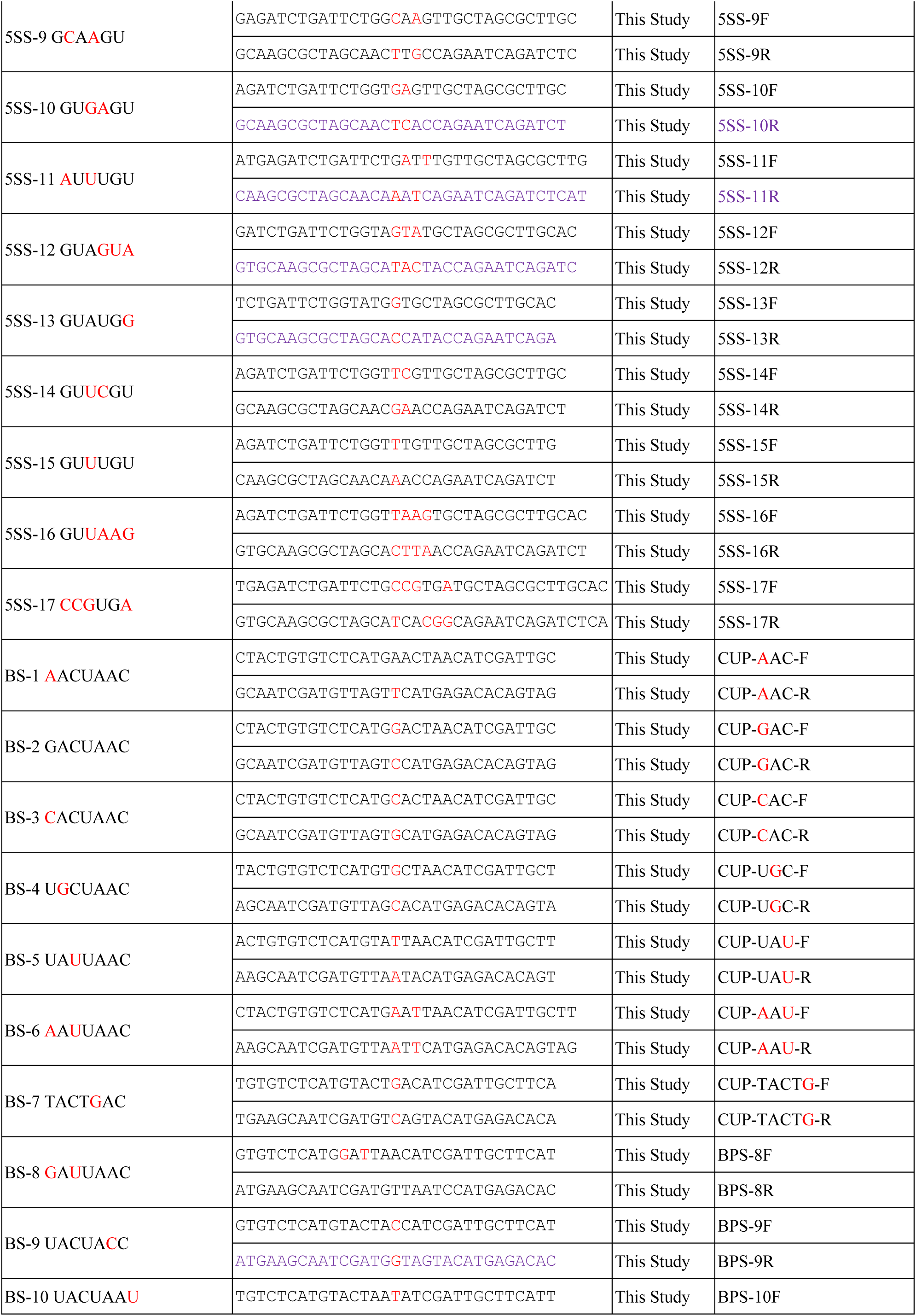
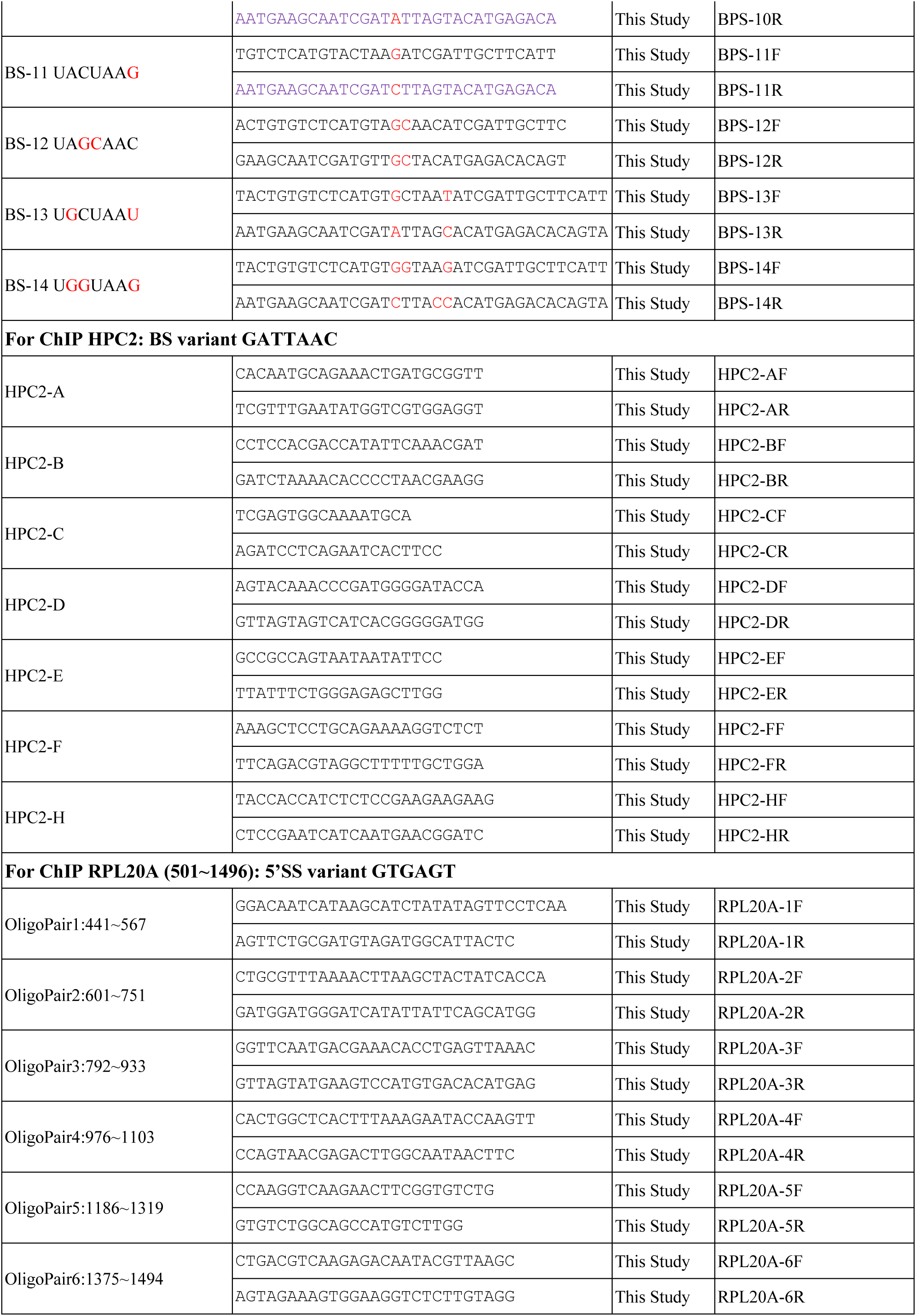
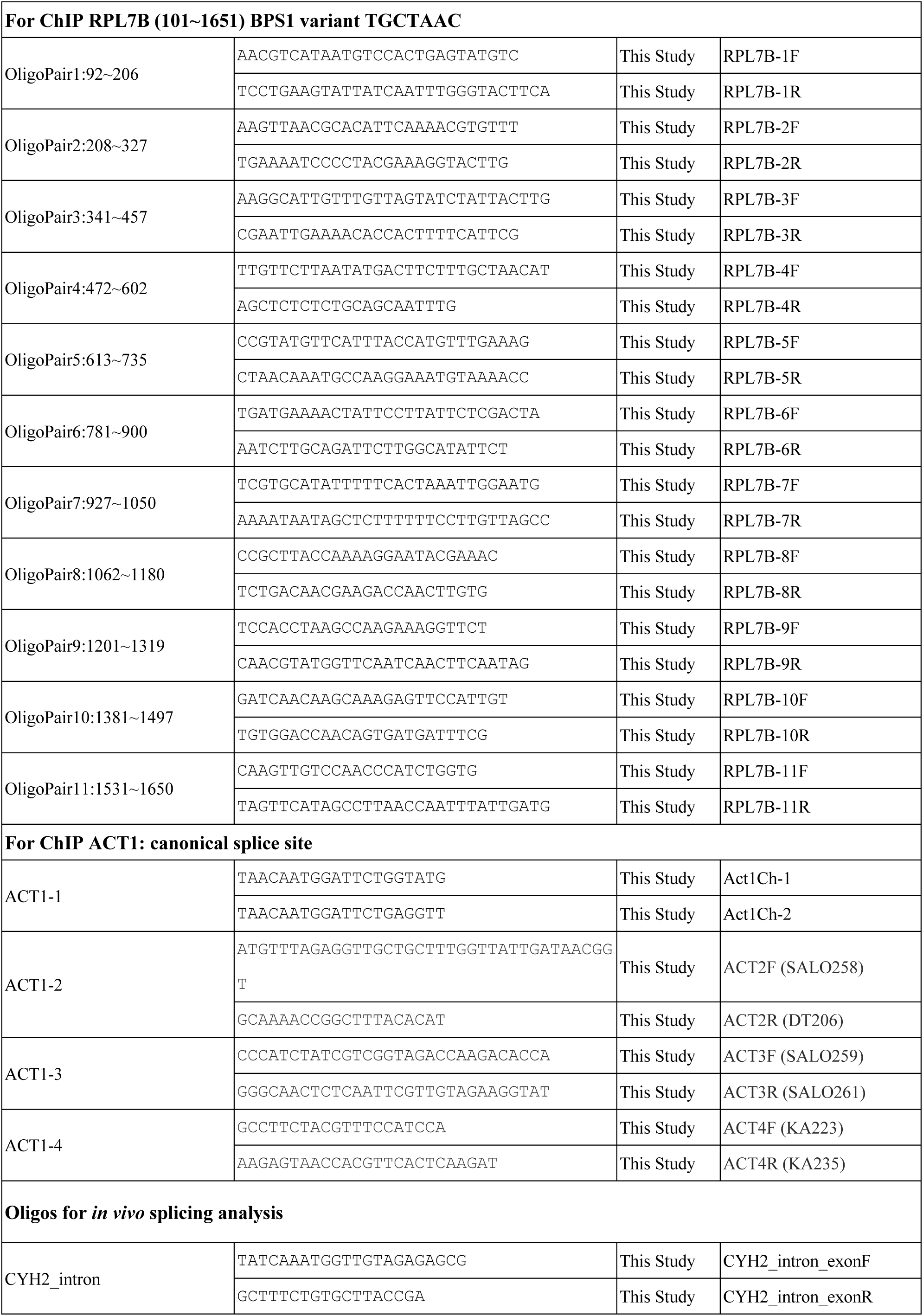
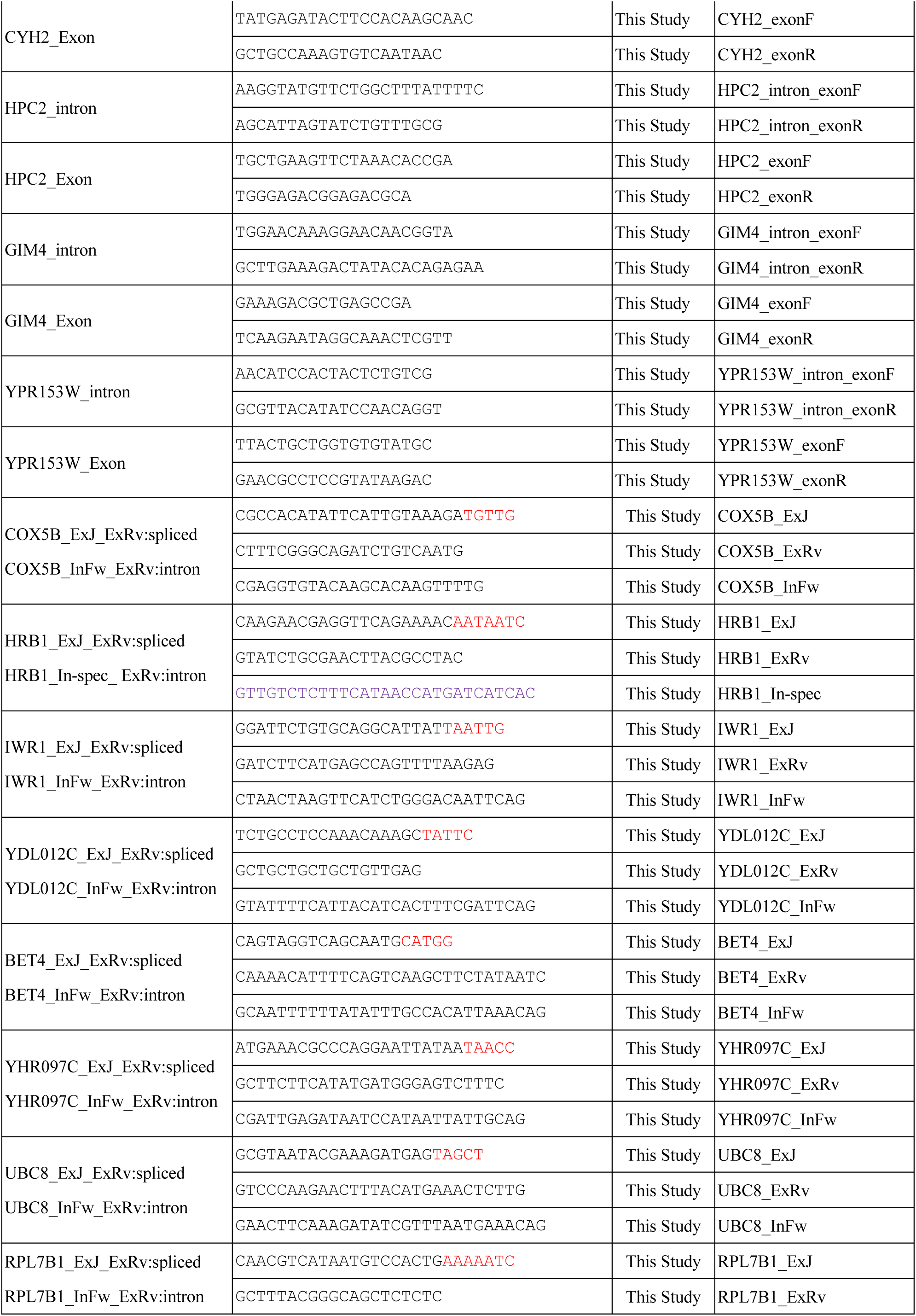
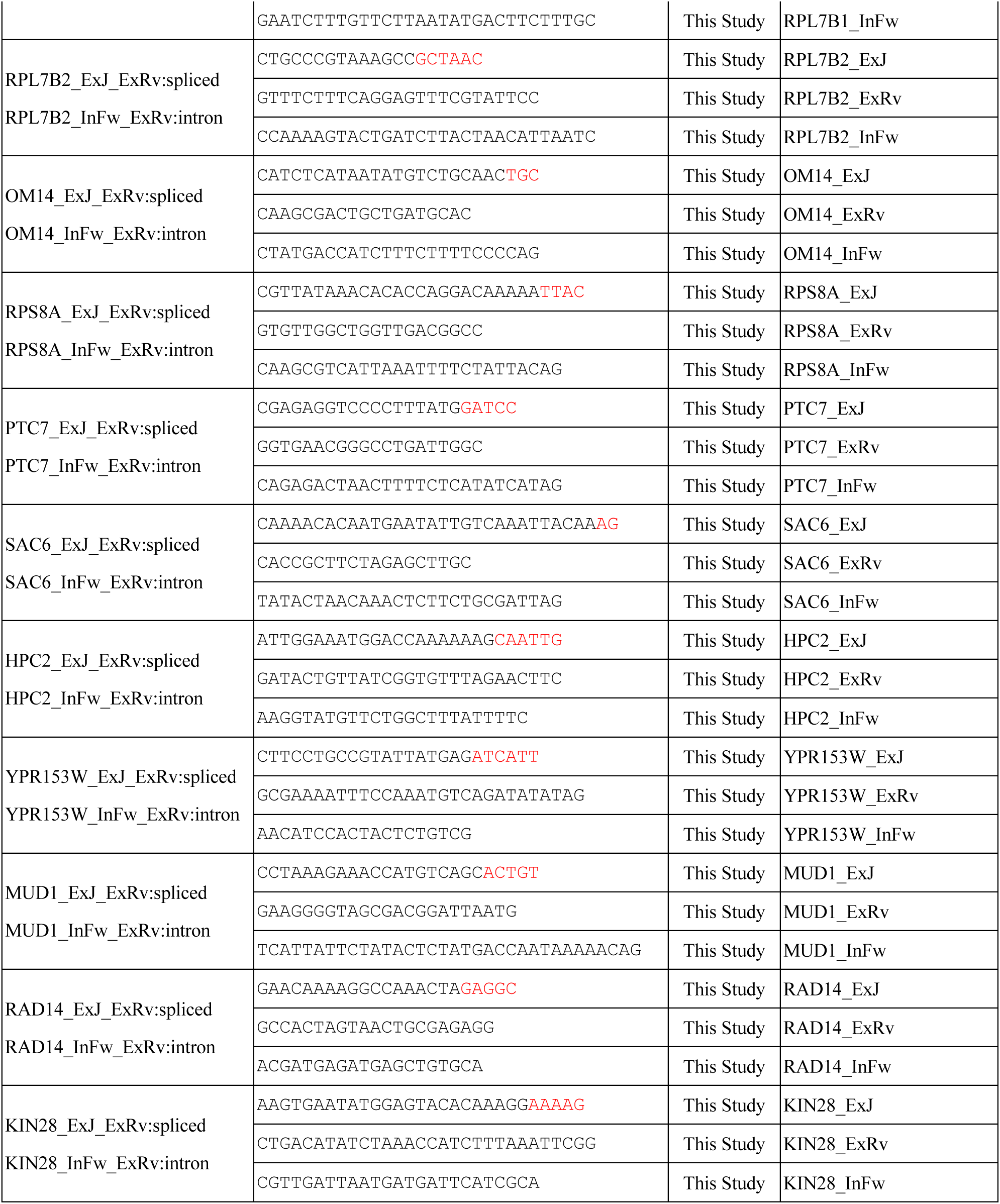
Oligonucleotide; related to STAR Methods.

## References

1. Galej, W.P., Nguyen, T.H.D., Newman, A.J., and Nagai, K. (2014). Structural studies of the spliceosome: zooming into the heart of the machine. Curr. Opin. Struct. Biol. 25, 57–66. 10.1016/j.sbi.2013.12.002.

2. Chen, W., and Moore, M.J. (2014). The spliceosome: disorder and dynamics defined. Curr. Opin. Struct. Biol. 24, 141–149. 10.1016/j.sbi.2014.01.009.

3. Chang, T.-H., Tung, L., Yeh, F.-L., Chen, J.-H., and Chang, S.-L. (2013). Functions of the DExD/H-box proteins in nuclear pre-mRNA splicing. Biochim. Biophys. Acta BBA - Gene Regul. Mech. 1829, 764–774. 10.1016/j.bbagrm.2013.02.006.

4. Wahl, M.C., Will, C.L., and Lührmann, R. (2009). The spliceosome: design principles of a dynamic RNP machine. Cell 136, 701–718. 10.1016/j.cell.2009.02.009.

5. Lacadie, S.A., Tardiff, D.F., Kadener, S., and Rosbash, M. (2006). In vivo commitment to yeast cotranscriptional splicing is sensitive to transcription elongation mutants. Genes Dev. 20, 2055–2066. 10.1101/gad.1434706.

6. Shcherbakova, I., Hoskins, A.A., Friedman, L.J., Serebrov, V., Corrêa, I.R., Xu, M.-Q., Gelles, J., and Moore, M.J. (2013). Alternative spliceosome assembly pathways revealed by single-molecule fluorescence microscopy. Cell Rep. 5, 151–165. 10.1016/j.celrep.2013.08.026.

7. Neuvéglise, C., Marck, C., and Gaillardin, C. (2011). The intronome of budding yeasts. C. R. Biol. 334, 662–670. 10.1016/j.crvi.2011.05.015.

8. Kao, H.-Y., and Siliciano, P.G. (1996). Identification of Prp40, a novel essential yeast splicing factor associated with the U1 small nuclear ribonucleoprotein particle. Mol. Cell. Biol. 16, 960–967. 10.1128/MCB.16.3.960.

9. Murphy, M.W., Olson, B.L., and Siliciano, P.G. (2004). The yeast splicing factor Prp40p contains functional leucine-rich nuclear export signals that are essential for splicing. Genetics 166, 53–65. 10.1534/genetics.166.1.53.

10. Blum, M., Andreeva, A., Florentino, L.C., Chuguransky, S.R., Grego, T., Hobbs, E., Pinto, B.L., Orr, A., Paysan-Lafosse, T., Ponamareva, I., et al. (2025). InterPro: the protein sequence classification resource in 2025. Nucleic Acids Res. 53, D444–D456. 10.1093/nar/gkae1082.

11. Abovich, N., and Rosbash, M. (1997). Cross-intron bridging interactions in the yeast commitment complex are conserved in mammals. Cell 89, 403–412. 10.1016/S0092-8674(00)80221-4.

12. Wiesner, S., Stier, G., Sattler, M., and Macias, M.J. (2002). Solution structure and ligand recognition of the WW domain pair of the yeast splicing factor Prp40. J. Mol. Biol. 324, 807–822. 10.1016/S0022-2836(02)01145-2.

13. Salah, Z., Alian, A., and Aqeilan, R.I. (2012). WW domain-containing proteins: retrospectives and the future. Front. Biosci.-Landmark 17, 331–348. 10.2741/3930.

14. Lapi, E., Di Agostino, S., Donzelli, S., Gal, H., Domany, E., Rechavi, G., Pandolfi, P.P., Givol, D., Strano, S., Lu, X., et al. (2008). PML, YAP, and p73 are components of a proapoptotic autoregulatory feedback loop. Mol. Cell 32, 803–814. 10.1016/j.molcel.2008.11.019.

15. Aqeilan, R.I., Pekarsky, Y., Herrero, J.J., Palamarchuk, A., Letofsky, J., Druck, T., Trapasso, F., Han, S.-Y., Melino, G., Huebner, K., et al. (2004). Functional association between Wwox tumor suppressor protein and p73, a p53 homolog. Proc. Natl. Acad. Sci. 101, 4401–4406. 10.1073/pnas.0400805101.

16. Becerra, S., Montes, M., Hernández-Munain, C., and Suñé, C. (2015). Prp40 pre-mRNA processing factor 40 homolog B (PRPF40B) associates with SF1 and U2AF65 and modulates alternative pre-mRNA splicing in vivo. RNA 21, 438–457. 10.1261/rna.047258.114.

17. Lin, K.-T., Lu, R.-M., and Tarn, W.-Y. (2004). The WW domain-containing proteins interact with the early spliceosome and participate in pre-mRNA splicing in vivo. Mol. Cell. Biol. 24, 9176–9185. 10.1128/MCB.24.20.9176-9185.2004.

18. Huibregtse, J.M., Scheffner, M., Beaudenon, S., and Howley, P.M. (1995). A family of proteins structurally and functionally related to the E6-AP ubiquitin-protein ligase. Proc. Natl. Acad. Sci. 92, 2563–2567. 10.1073/pnas.92.7.2563.

19. Aqeilan, R.I., Trapasso, F., Hussain, S., Costinean, S., Marshall, D., Pekarsky, Y., Hagan, J.P., Zanesi, N., Kaou, M., Stein, G.S., et al. (2007). Targeted deletion of Wwox reveals a tumor suppressor function. Proc. Natl. Acad. Sci. 104, 3949–3954. 10.1073/pnas.0609783104.

20. Paige, A.J.W., Taylor, K.J., Taylor, C., Hillier, S.G., Farrington, S., Scott, D., Porteous, D.J., Smyth, J.F., Gabra, H., and Watson, J.E.V. (2001). WWOX: A candidate tumor suppressor gene involved in multiple tumor types. Proc. Natl. Acad. Sci. 98, 11417–11422. 10.1073/pnas.191175898.

21. Hansson, J.H., Schild, L., Lu, Y., Wilson, T.A., Gautschi, I., Shimkets, R., Nelson-Williams, C., Rossier, B.C., and Lifton, R.P. (1995). A de novo missense mutation of the beta subunit of the epithelial sodium channel causes hypertension and Liddle syndrome, identifying a proline-rich segment critical for regulation of channel activity. Proc. Natl. Acad. Sci. 92, 11495–11499. 10.1073/pnas.92.25.11495.

22. Faber, P.W., Barnes, G.T., Srinidhi, J., Chen, J., Gusella, J.F., and MacDonald, M.E. (1998). Huntingtin interacts with a family of WW domain proteins. Hum. Mol. Genet. 7, 1463–1474. 10.1093/hmg/7.9.1463.

23. Passani, L.A., Bedford, M.T., Faber, P.W., McGinnis, K.M., Sharp, A.H., Gusella, J.F., Vonsattel, J.-P., and MacDonald, M.E. (2000). Huntingtin’s WW domain partners in Huntington’s disease post-mortem brain fulfill genetic criteria for direct involvement in Huntington’s disease pathogenesis. Hum. Mol. Genet. 9, 2175–2182. 10.1093/hmg/9.14.2175.

24. Buschdorf, J.P., and Strätling, W.H. (2004). A WW domain binding region in methyl-CpG-binding protein MeCP2: impact on Rett syndrome. J. Mol. Med. 82, 135–143. 10.1007/s00109-003-0497-9.

25. Papaemmanuil, E., Cazzola, M., Boultwood, J., Malcovati, L., Vyas, P., Bowen, D., Pellagatti, A., Wainscoat, J.S., Hellstrom-Lindberg, E., Gambacorti-Passerini, C., et al. (2011). Somatic SF3B1 mutation in myelodysplasia with ring sideroblasts. N. Engl. J. Med. 365, 1384–1395. 10.1056/NEJMoa1103283.

26. Yoshida, K., Sanada, M., Shiraishi, Y., Nowak, D., Nagata, Y., Yamamoto, R., Sato, Y., Sato-Otsubo, A., Kon, A., Nagasaki, M., et al. (2011). Frequent pathway mutations of splicing machinery in myelodysplasia. Nature 478, 64–69. 10.1038/nature10496.

27. Görnemann, J., Barrandon, C., Hujer, K., Rutz, B., Rigaut, G., Kotovic, K.M., Faux, C., Neugebauer, K.M., and Séraphin, B. (2011). Cotranscriptional spliceosome assembly and splicing are independent of the Prp40p WW domain. RNA 17, 2119–2129. 10.1261/rna.02646811.

28. Li, X., Liu, S., Zhang, L., Issaian, A., Hill, R.C., Espinosa, S., Shi, S., Cui, Y., Kappel, K., Das, R., et al. (2019). A unified mechanism for intron and exon definition and back-splicing. Nature 573, 375–380. 10.1038/s41586-019-1523-6.

29. Zhang, Z., Rigo, N., Dybkov, O., Fourmann, J.-B., Will, C.L., Kumar, V., Urlaub, H., Stark, H., and Lührmann, R. (2021). Structural insights into how Prp5 proofreads the pre-mRNA branch site. Nature 596, 296–300. 10.1038/s41586-021-03789-5.

30. Abramson, J., Adler, J., Dunger, J., Evans, R., Green, T., Pritzel, A., Ronneberger, O., Willmore, L., Ballard, A.J., Bambrick, J., et al. (2024). Accurate structure prediction of biomolecular interactions with AlphaFold 3. Nature 630, 493–500. 10.1038/s41586-024-07487-w.

31. Ester, C., and Uetz, P. (2008). The FF domains of yeast U1 snRNP protein Prp40 mediate interactions with Luc7 and Snu71. BMC Biochem. 9, 29. 10.1186/1471-2091-9-29.

32. Martínez-Lumbreras, S., Träger, L.K., Mulorz, M.M., Payr, M., Dikaya, V., Hipp, C., König, J., and Sattler, M. (2024). Intramolecular autoinhibition regulates the selectivity of PRPF40A tandem WW domains for proline-rich motifs. Nat. Commun. 15, 3888. 10.1038/s41467-024-48004-x.

33. Tong, A.H.Y., Evangelista, M., Parsons, A.B., Xu, H., Bader, G.D., Pagé, N., Robinson, M., Raghibizadeh, S., Hogue, C.W.V., Bussey, H., et al. (2001). Systematic genetic analysis with ordered arrays of yeast deletion mutants. Science 294, 2364–2368. 10.1126/science.1065810.

34. Tong, A.H.Y., and Boone, C. (2006). Synthetic genetic array analysis in Saccharomyces cerevisiae. In Yeast Protocol Methods in Molecular Biology., W. Xiao, ed. (Humana Press), pp. 171–191. 10.1385/1-59259-958-3:171.

35. Kistler, A.L., and Guthrie, C. (2001). Deletion of MUD2, the yeast homolog of U2AF65, can bypass the requirement for Sub2, an essential spliceosomal ATPase. Genes Dev. 15, 42–49. 10.1101/gad.851601.

36. Caspary, F., and Séraphin, B. (1998). The yeast U2A′/U2B″ complex is required for pre-spliceosome formation. EMBO J. 17, 6348–6358. 10.1093/emboj/17.21.6348.

37. Plaschka, C., Lin, P.-C., Charenton, C., and Nagai, K. (2018). Prespliceosome structure provides insights into spliceosome assembly and regulation. Nature 559, 419–422. 10.1038/s41586-018-0323-8.

38. Qiu, Z.R., Chico, L., Chang, J., Shuman, S., and Schwer, B. (2012). Genetic interactions of hypomorphic mutations in the m7G cap-binding pocket of yeast nuclear cap binding complex: an essential role for Cbc2 in meiosis via splicing of MER3 pre-mRNA. RNA 18, 1996–2011. 10.1261/rna.033746.112.

39. Pleiss, J.A., Whitworth, G.B., Bergkessel, M., and Guthrie, C. (2007). Transcript specificity in yeast pre-mRNA splicing revealed by mutations in core spliceosomal components. PLOS Biol. 5, e90. 10.1371/journal.pbio.0050090.

40. Inada, M., and Pleiss, J.A. (2010). Chapter 3 - Genome-wide approaches to monitor pre-mRNA splicing. In Methods in Enzymology Guide to Yeast Genetics: Functional Genomics, Proteomics, and Other Systems Analysis. (Academic Press), pp. 51–75. 10.1016/S0076-6879(10)70003-3.

41. Juneau, K., Palm, C., Miranda, M., and Davis, R.W. (2007). High-density yeast-tiling array reveals previously undiscovered introns and extensive regulation of meiotic splicing. Proc. Natl. Acad. Sci. 104, 1522–1527. 10.1073/pnas.0610354104.

42. Spingola, M., Grate, L., Haussler, D., and Ares, M. (1999). Genome-wide bioinformatic and molecular analysis of introns in Saccharomyces cerevisiae. RNA 5, 221–234.

43. Engel, S.R., Aleksander, S., Nash, R.S., Wong, E.D., Weng, S., Miyasato, S.R., Sherlock, G., and Cherry, J.M. (2025). Saccharomyces Genome Database: advances in genome annotation, expanded biochemical pathways, and other key enhancements. Genetics 229, iyae185. 10.1093/genetics/iyae185.

44. Lesser, C.F., and Guthrie, C. (1993). Mutational analysis of pre-mRNA splicing in Saccharomyces cerevisiae using a sensitive new reporter gene, CUP1. Genetics 133, 851–863. 10.1093/genetics/133.4.851.

45. Chalivendra, S., Shi, S., Li, X., Kuang, Z., Giovinazzo, J., Zhang, L., Rossi, J., Wang, J., Saviola, A.J., Welty, R., et al. (2024). Selected humanization of yeast U1 snRNP leads to global suppression of pre-mRNA splicing and mitochondrial dysfunction in the budding yeast. RNA 30, 1070–1088. 10.1261/rna.079917.123.

46. Lipp, J.J., Marvin, M.C., Shokat, K.M., and Guthrie, C. (2015). SR protein kinases promote splicing of nonconsensus introns. Nat. Struct. Mol. Biol. 22, 611–617. 10.1038/nsmb.3057.

47. Goldstrohm, A.C., Albrecht, T.R., Suñé, C., Bedford, M.T., and Garcia-Blanco, M.A. (2001). The transcription elongation factor CA150 interacts with RNA polymerase II and the pre-mRNA splicing factor SF1. Mol. Cell. Biol. 21, 7617–7628. 10.1128/MCB.21.22.7617-7628.2001.

48. Dittmar, J.C., Reid, R.J., and Rothstein, R. (2010). ScreenMill: A freely available software suite for growth measurement, analysis and visualization of high-throughput screen data. BMC Bioinformatics 11, 353. 10.1186/1471-2105-11-353.

49. Pleiss, J.A., Whitworth, G.B., Bergkessel, M., and Guthrie, C. (2007). Rapid, transcript-specific changes in splicing in response to environmental stress. Mol. Cell 27, 928–937. 10.1016/j.molcel.2007.07.018.

50. Chen, J.Y.-F., Stands, L., Staley, J.P., Jackups, R.R., Latus, L.J., and Chang, T.-H. (2001). Specific alterations of U1-C protein or U1 small nuclear RNA can eliminate the requirement of Prp28p, an essential DEAD box splicing factor. Mol. Cell 7, 227–232. 10.1016/S1097-2765(01)00170-8.

51. Lacadie, S.A., and Rosbash, M. (2005). Cotranscriptional spliceosome assembly dynamics and the role of U1 snRNA:5′ss base pairing in yeast. Mol. Cell 19, 65–75. 10.1016/j.molcel.2005.05.006.

52. Meng, E.C., Goddard, T.D., Pettersen, E.F., Couch, G.S., Pearson, Z.J., Morris, J.H., and Ferrin, T.E. (2023). UCSF ChimeraX: Tools for structure building and analysis. Protein Sci. 32, e4792. 10.1002/pro.4792.

## Supplemental References

1. Inada, M., and Pleiss, J.A. (2010). Chapter 3 - Genome-wide approaches to monitor pre-mRNA splicing. In Methods in Enzymology Guide to Yeast Genetics: Functional Genomics, Proteomics, and Other Systems Analysis. (Academic Press), pp. 51–75. 10.1016/S0076-6879(10)70003-3.

2. Pleiss, J.A., Whitworth, G.B., Bergkessel, M., and Guthrie, C. (2007). Rapid, transcript-specific changes in splicing in response to environmental stress. Mol. Cell 27, 928–937. 10.1016/j.molcel.2007.07.018.

3. Pleiss, J.A., Whitworth, G.B., Bergkessel, M., and Guthrie, C. (2007). Transcript specificity in yeast pre-mRNA splicing revealed by mutations in core spliceosomal components. PLOS Biol. 5, e90. 10.1371/journal.pbio.0050090.

4. Abramson, J., Adler, J., Dunger, J., Evans, R., Green, T., Pritzel, A., Ronneberger, O., Willmore, L., Ballard, A.J., Bambrick, J., et al. (2024). Accurate structure prediction of biomolecular interactions with AlphaFold 3. Nature 630, 493–500. 10.1038/s41586-024-07487-w.

5. Zhang, Z., Rigo, N., Dybkov, O., Fourmann, J.-B., Will, C.L., Kumar, V., Urlaub, H., Stark, H., and Lührmann, R. (2021). Structural insights into how Prp5 proofreads the pre-mRNA branch site. Nature 596, 296–300. 10.1038/s41586-021-03789-5.

6. Wiesner, S., Stier, G., Sattler, M., and Macias, M.J. (2002). Solution structure and ligand recognition of the WW domain pair of the yeast splicing factor Prp40. J. Mol. Biol. 324, 807–822. 10.1016/S0022-2836(02)01145-2.

7. Lipp, J.J., Marvin, M.C., Shokat, K.M., and Guthrie, C. (2015). SR protein kinases promote splicing of nonconsensus introns. Nat. Struct. Mol. Biol. 22, 611–617. 10.1038/nsmb.3057.

